# Ligand supplementation restores the cancer therapy efficacy of an antirheumatic drug auranofin from serum inactivation

**DOI:** 10.1101/2024.01.25.577173

**Authors:** Yuan Wang, Bei Cao, Qianqian Wang, Xin Fang, Junjian Wang, Albert S. C. Chan, Xiaolin Xiong, Taotao Zou

## Abstract

Auranofin, an FDA-approved antirheumatic gold drug, has gained ongoing interest in clinical studies for treating advanced or recurrent tumors. However, gold ion’s dynamic thiol exchange nature strongly attenuates its bioactivity due to the fast formation of covalent albumin-gold adducts. Here we report that newly-added thiols can modulate the dynamic albumin-gold binding and recover the therapeutic efficacy. Initially, we found that auranofin supplemented with its own thiol ligand, TGTA (1-thio-β-D-glucose tetraacetate), significantly restored the anticancer activities in cells and patient-derived xenograft models. Then, screening a collection of ligand fragments followed by machine learning evaluation unveiled diverse synergizing thiols, including pantethine that effectuates auranofin at a low dosage used for rheumatoid arthritis. Interestingly, the thiol exchange inside cells accounts for a cuproptosis-like phenotype induced by auranofin. Together, we believe the ligand-enabled dynamic modulation strategy is of value to researchers and clinicians contemplating metallodrugs and ligand-like molecules in cancer therapy.

---

As representative anticancer agents, platinum drugs were taken by 10-20% of all cancer patients (https://www.cancer.gov/research/progress/discovery/cisplatin). Their therapeutic value continues to be high in the clinic and clinical trials as synergistic agents with other cutting-edge cancer therapies^1–6^. The potent bioactivities underlying platinum drugs are associated with their distinctive ligand exchange kinetics before and after entering cancer cells. In blood, cisplatin, for example, displays fair stability against hydrolysis and off-target binding due to the concentrated chloride, but once inside cancer cells, it undergoes dynamic ligand hydrolysis and exchange to end up with binding biomolecular targets^7–9^. Unfortunately, such a kinetic trait seems hardly endowed by other metal compounds^10^, and no metal anticancer drugs other than platinum have been approved so far^11^. Extensive efforts have been made on the stability-reactivity trade-off of metallodrugs based on structure-activity-relationship in past decades^1,11–14^ but with limited success.

In addition to designing new molecules, repurposing previously approved metallodrugs appears to be a more efficient and less costly way^15–19^. In this regard, auranofin, a US Food and Drug Administration (FDA)-approved gold drug for rheumatoid arthritis (RA), has undergone substantial preclinical and clinical studies for repurposing against advanced or recurrent cancers^20–23^. This rheumatic gold drug has a high binding affinity to cysteine- and selenocysteine-containing proteins and acts as a pro-oxidative agent to kill various types of cancer cells including the cisplatin-resistant variants^24–35^. Still, a similar challenge is that the dynamic thiol exchange (DyThex) nature of gold leads to instability once the drug encounters the off-target thiols before reaching the tumor. In particular, the most abundant serum protein albumin (∼40 mg/mL) with a free thiol group (Cys34) can displace the 1-thio-β-D-glucose tetraacetate (TGTA) ligand of auranofin to form a protein-gold adduct in a fast and complete manner^36–38^, which leads to poor bioavailability and unsatisfactory anticancer efficacy *in vivo*^39–41^.

Given that the albumin-auranofin binding is a “thiol-to-thiol” exchange reaction, we conceived the possibility of rescuing active gold by additionally introducing small thiol molecules to modulate the dynamic equilibrium. Herein we report that supplementing auranofin with TGTA enhanced the anticancer efficacy in both cell lines and animal models, including patient-derived tumor xenografts. Further, cell-based screening of >200 commercially available ligand fragments followed by a machine learning evaluation helped identify a group of new thiol molecules, particularly pantethine that can boost the anticancer activity of auranofin at a low dosage equal to those permitted for treating RA in humans. Interestingly, we also found that the intracellular DyThex enables auranofin to engage in a cuproptosis-like mode of action.

## Results

### Thiol exchange with auranofin in the multidrug combo CUSP9v3

Some clinical trials in recent years have been associated with auranofin (Fig. 1a) for cancer therapy (https://classic.clinicaltrials.gov). Among them, the trial NCT02770378 drew our attention as it used 9 drugs (including auranofin) in combination with temozolomide for recurrent glioblastoma treatment^42,43^ (Fig. 1b). Each drug is selected for targeting a certain pathway with a synergistic effect against the resistance mechanisms of this cancer. The phase Ib/IIa experiment has confirmed the safety of this treatment regime (CUP9v3) and a randomized phase II trial has also been completed.

**Fig. 1.**
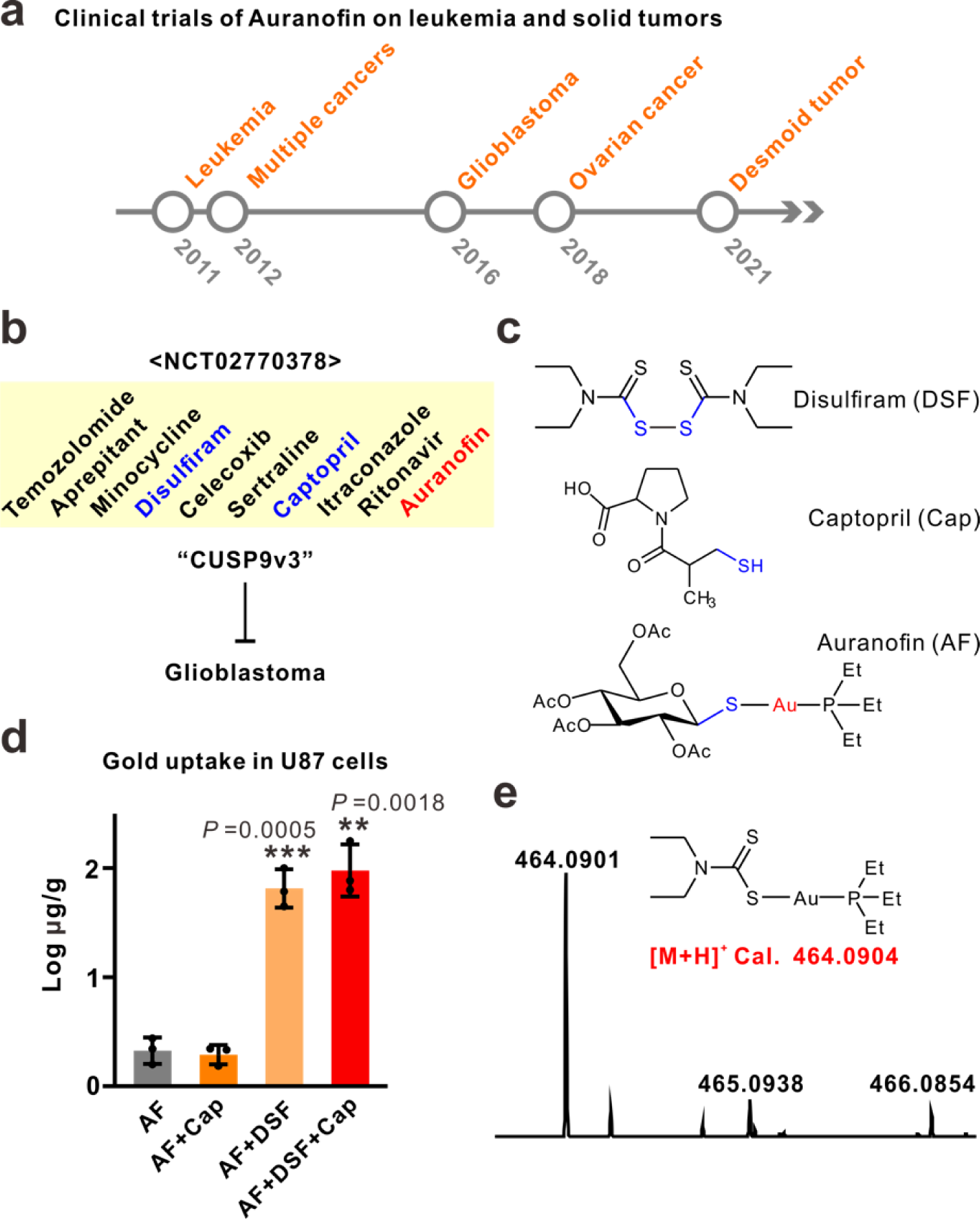
Thiol exchange in auranofin-related clinical trials. **a,** a timeline summary of auranofin-repurposing clinical trials on cancer therapy in recent years. **b,** drugs used in clinical trial NCT02770378 for glioblastoma. **c,** the chemical structures of indicated drugs. **d,** cellular gold uptake affected by the cotreatment with thiol-containing drugs in U87 cells. 3 μM of auranofin was used in all groups. Data are shown as mean ± s. d. of three independent experiments. Significance was calculated by unpaired, two-tailed *t*-test. **e,** mass spectrometry identified the formation of a DEDT-gold-phosphine molecule after an *in vitro* incubation of auranofin, captopril, and disulfiram. The spectrum was extracted from 463.0000 to 467.0000 *m/z* [M+H]^+^ values with the predicted *m/z* labeled in red. The result is representative of three independent experiments.

Besides auranofin, two thiol-containing drugs disulfiram and captopril (Fig. 1c), which are in large molar excess (190- and 260-fold as compared to auranofin, respectively), were also included in CUP9v3. To check their possible impacts on auranofin, we first assessed the intracellular gold content after a short period (10 min) of the co-treatments on a glioblastoma cell line U87 following their ratios in CUP9v3. The treatments were conducted in the presence of 30% FBS to mimic the physiological interference from albumins. Interestingly, the addition of disulfiram largely enhanced the cellular uptake of gold by over 50-fold at this serum-containing condition (Fig. 1d). Captopril, however, had no direct effect on the uptake. Incubating the three drugs in aqueous solution generated a new compound in which diethyldithiocarbamate (DEDT), the reduced product of disulfiram, displaced the TGTA ligand in auranofin (Fig. 1e). Thus, the drug interactions occur in this multidrug combo that may contribute to the bioavailability of gold.

### Albumin blockade on auranofin is reversed by excess TGTA ligand

In CUP9v3, the reduced DEDT is at large excess, so we considered that it may compete with albumin, impeding the formation of cell-impermeable protein-gold complex and improving the bioavailability of gold. However, DEDT has been known to inhibit NPL4 and cause strong cytotoxicity^44^. Therefore, we sought to harness TGTA, a ligand with minimal cytotoxicity (IC_50_ > 300 μM), to further support our hypothesis that excess TGTA ligand may shift the equilibrium from the albumin-gold adduct back to auranofin and recover cytotoxicity (Fig. 2a). To begin with, we monitored the reaction between albumin and auranofin by HPLC. As expected, the peak of auranofin (retention time 3.78 min) disappeared when mixed with albumin (Fig. 2b), in association with the formation of an albumin-Au-PEt_3_ adduct (characterized by HRMS, Extended Data fig. 1). Of note, the peak of auranofin reappeared in the LC chromatogram after adding excess TGTA into the albumin-auranofin mixture (Fig. 2b). Encouraged by this phenomenon, we tested the effects of TGTA at the cellular level. Interestingly, the gold uptake was significantly reduced with increasing concentration of serum, but recovered by supplementing with TGTA (Fig. 2c). Next, we looked into the effects of adding TGTA on the cytotoxicity. While the cytotoxicity of auranofin was largely suppressed by albumins in the serum (Extended Data Fig. 2), a TGTA dose-dependent recovery was found with a fixed concentration of auranofin (Fig. 2d). As a control, the glucose pentaacetate, a thiol-free homolog of TGTA, did not influence the cytotoxicity (Fig. 2d). Consistent with increased cellular gold contents and cytotoxicity, the activity of thioredoxin reductase (TrxR), a primary target of auranofin^45–47^, was significantly inhibited by the additional TGTA ligands in the living HCT116 cells (Fig. 2e).

**Fig. 2.**
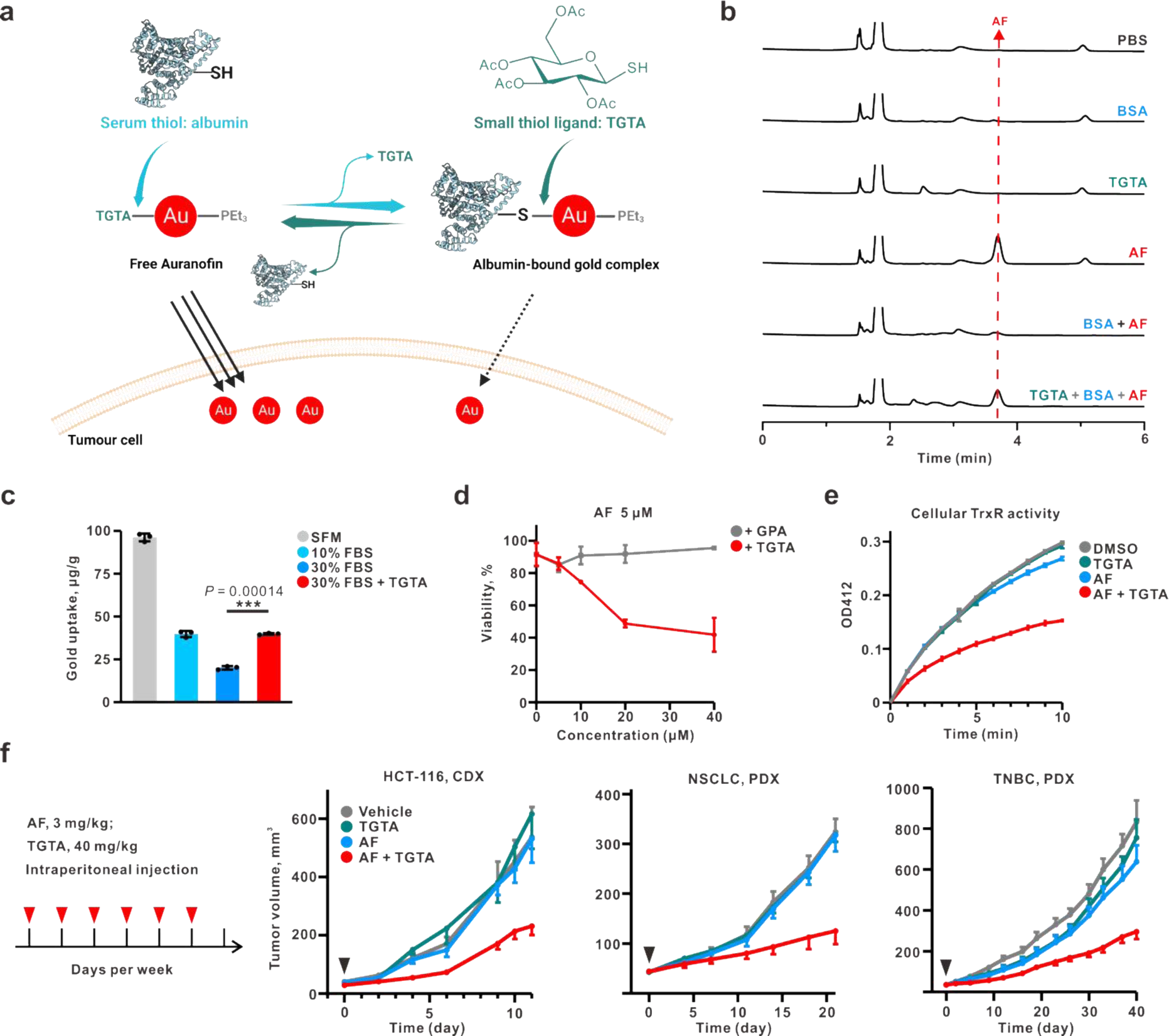
TGTA restores the anticancer activity of albumin-bound auranofin. **a,** a scheme for harnessing the dynamic thiol exchange to restore anticancer efficacy from albumin-bound gold. TGTA, thioglucose tetraacetate. **b,** HPLC analysis at 214 nm on the thiol exchange reaction *in vitro*. The peak of auranofin was marked by a dotted red line. **c,** cellular gold uptake of HCT116 cells under indicated conditions of FBS. 3 μM of auranofin was used in all groups. The values were normalized by the protein amount extracted. Data are shown as mean ± s. d. of three independent experiments. Significance was calculated by unpaired, two-tailed *t*-test. **d,** dose-dependent cytotoxicity boosting of auranofin by glucose homologs. GPA, Glucose pentaacetate. **e,** measuring cellular thioredoxin reductase (TrxR) activity using HCT116 upon indicated treatments. For **d-e,** mean ± s. d., 3 biological replicates. **f,** mouse models to evaluate the *in vivo* tumor suppression of auranofin. Drugs were injected intraperitoneally 6 times per week and the initial administration day was marked by black arrows. An HCT116 xenograft and two patient-derived xenografts (non-small cell lung carcinoma triple-negative breast cancer) were conducted. HCT116 tumors: Vehicle, n=7 mice; TGTA, n=6 mice; AF, n=6 mice; AF+TGTA, n=9 mice. NSCLC PDX tumors: Vehicle, n = 7 mice; TGTA, n = 6 mice; AF, n = 6 mice; AF+TGTA, n = 7 mice. TNBC PDX tumors: Vehicle, n = 9 mice; TGTA, n = 9 mice; AF, n = 8 mice; AF+TGTA, n = 8 mice. Data were shown as mean ± s. e. m.

Next, we examined the efficacy of the “auranofin + TGTA” combo in mouse models (Fig. 2f). An early study claimed that only P388 leukemia, out of 15 tumor mouse models (blood and solid tumors), was sensitive to auranofin treatment at a dose of 12 mg/kg intraperitoneally^40^. Later works showed that 10 mg/kg of auranofin was normally required to achieve the anticancer effect in murine models^46,48–50^. Here we used 3 mg/kg, a dosage with minimal anti-tumor activity *in vivo* in auranofin treatment alone. In the mice-bearing HCT116 xenograft, TGTA (40 mg/kg) significantly enhanced the tumor suppression of auranofin compared to vehicle control after intraperitoneal injection, but TGTA or auranofin alone failed to suppress tumor growth. Then we sought to evaluate the responses in two patient-derived xenograft models, non-small cell lung carcinoma (NSCLC) and triple-negative breast cancer (TNBC), respectively. Similar results were observed in these two PDX models (Fig. 2f). To further check the impact of our combo on mouse physiology, we monitored the body weight of each mouse. In all three models, we found no mouse death or noticeable change in mouse body weight (Extended Data Fig. 3a). Besides, no pathological features were observed by inspecting internal organs (heart, kidney, liver, lung, spleen) in the mice of NSCLC PDX models (Extended Data Fig. 3b). These results suggest that TGTA can potentiate the antitumor activity of auranofin without apparent toxicity.

### Ligand screening unveils new molecules able to synergize auranofin

The successful use of TGTA drove us to mine more ligands with synergistic activities. A collection of 252 commercially available ligand fragments containing N, O, S, or P binding sites were screened at a constant concentration (50 μM) for their synergistic effect on auranofin. The cell viability of the combo group versus auranofin alone (“A+L”/A) was plotted as the effectiveness ratio (Fig. 3a). We selected those with a ratio less than 0.3 as the primary pool of active candidates and then tested their own cytotoxicity (Fig. 3b). Finally, 14 compounds with synergistic effects but also negligible cytotoxicity (<10 %) were picked as the “effective” ones. Notably, thiol-(including disulfide-) or phosphine-containing ones were significantly enriched in the effective pool (Fig. 3c), in line with the high affinity of gold ions towards these two types of ligands. Computational calculation using density functional theory (DFT) on different types of ligands (Cl, N, O, P, S) supports the screening outcomes as only thiol and phosphine ligands can lead to favorable shift of ligand exchange equilibrium (chemical equilibrium constant *K* close to 1 or higher, Extended Data Fig.4). For the thiols to be more physiologically relevant, we sought to compare the “effective” and “ineffective” thiols or disulfides. Based on the 354 calculated chemical descriptors for each thiol-containing molecule, we trained a random forest classification model to weigh their importance toward the effectiveness (Fig. 3d). As the results show, most descriptors related to molecular polarity and hydrophobicity were ranked top of all (Fig. 3d). This indicated that ligands with suitable lipophilicity can shuttle gold ions into the cells.

**Fig. 3.**
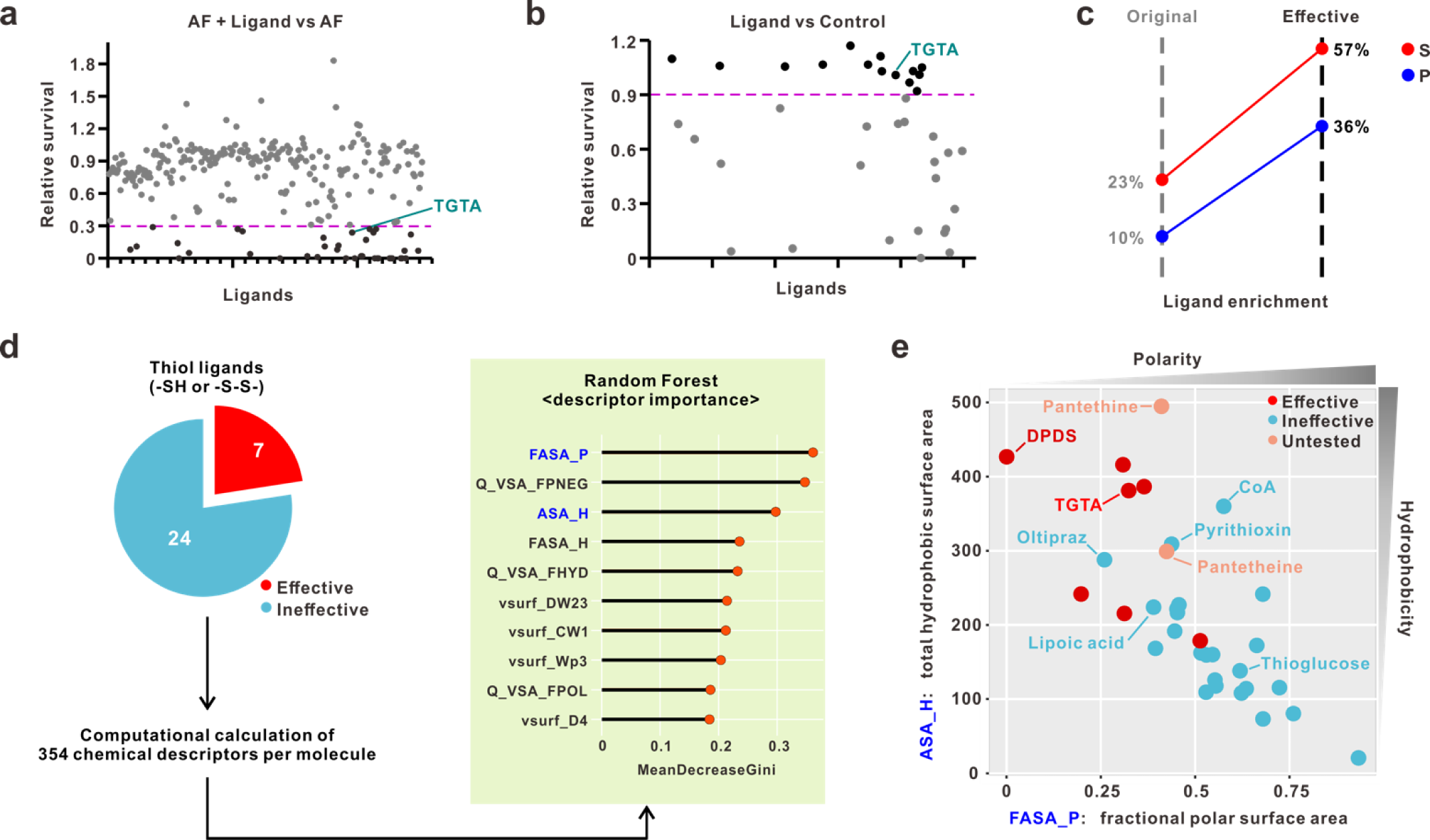
Ligand screening in synergy with auranofin. **a,** PC9 cell viability in response to cotreatments using 5 μM auranofin with 50 μM ligands from a chemical library containing 252 small molecules. Ligands with values under 0.3 (dotted line in pink) were picked for the next experiments. **b,** a counter-screening on the self-toxicity of the picked ligands. Ligands with less than 10% toxicity were picked as the effective synergistic ones. **c,** percentages of sulfur- (S) or phospho- (P) containing molecules in the original and effective ligands were compared. **d,** a workflow evaluating the synergy-associated chemical features using random forest classification based on 354 descriptors calculated by MOE software. The top 10 descriptors ranked by importance were listed with their MeanDecreaseGini values. **e,** plotting the thiol ligands by their values of two descriptors, FASA_P and ASA_H. Effective, ineffective, and untested molecules were colored as indicated. Names of some ligands were shown.

To visualize the classification, we plotted the thiol ligands using two top-ranking descriptors, FASA_P (fractional polar surface area) for polarity and ASA_H (total hydrophobic surface area) for hydrophobicity (Fig. 3e). According to the plot, effective ones were preferably located at the high hydrophobicity and low polarity region. Diphenyl disulfide (DPDS) with the most hydrophobicity indicated by the plot is a chemical reagent frequently used in organic synthesis. Noteworthily, a recent report claimed an anti-breast cancer activity of this compound in an apoptosis-promoting mechanism^51^. Oltipraz and pyrithioxin are ineffective ligands but are located near the region of the effective in the plot. Reasonably, these two agents still showed a potential to synergize auranofin in the primary screening with effectiveness ratios of 0.57 and 0.31, respectively (close to the set threshold of 0.3). Based on the analysis above, we hypothesized that chemical parameters related to hydrophobicity can serve as important criteria for choosing effective thiol ligands.

Next, we wondered whether hydrophilic thiols can shuttle gold ions to certain cell types. To this end, a further screening using hydrophilic but physiologically relevant thiols in a higher concentration on ten cell lines was performed (Fig. 4a). This comprises coenzyme A (CoA) ^52^, some of its derivatives, and other two thiols (lipoic acid, thioglucose). However, CoA, lipoic acid, and thioglucose were still less effective, although a relatively high auranofin-boosting activity was seen by CoA in Jurkat T cells. Interestingly, pantethine and its reduced monomeric form pantetheine came up with the strongest synergistic effects.

**Fig. 4.**
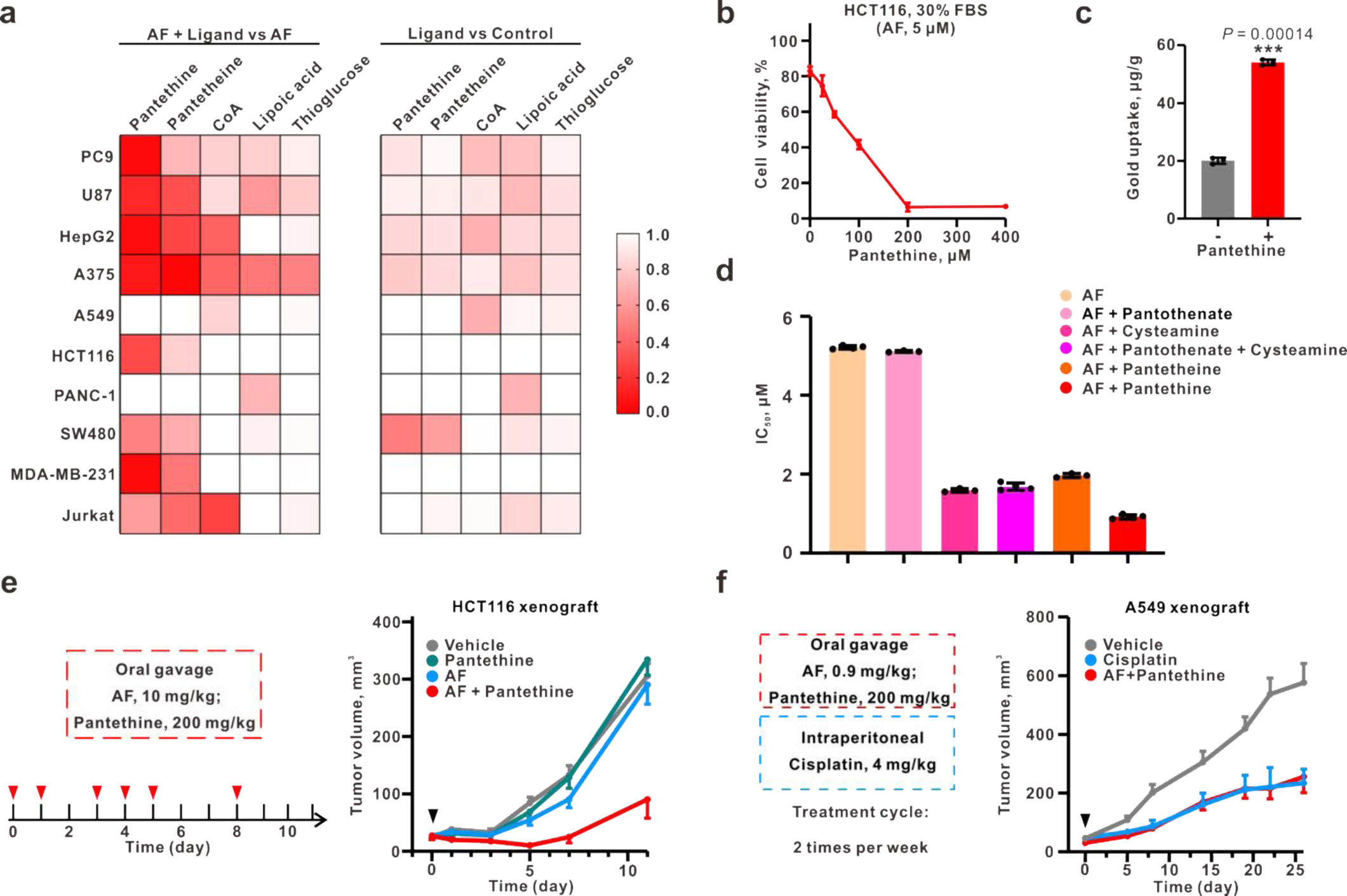
Synergistic effect of pantethine on auranofin. **a,** a small-scale cell line profiling using indicated agents. The color bar represents the mean value of relative survival in two independent experiments. **b,** the dose-dependent synergy of pantethine with auranofin. **c,** cellular gold uptake of HCT116 cells under 3 μM auranofin with or without 200 μM pantethine. The values were normalized by the protein amount extracted. Data are shown as mean ± s. d. of three independent experiments. Significance was calculated by unpaired, two-tailed *t*-test. **d,** the comparison of the synergistic activities with auranofin endowed by different pantothenate derivatives using HCT116 cells. Data are shown as mean ± s. d., 3 biologically independent experiments. Significance was calculated by unpaired, two-tailed *t*-test. **e,** mouse models using oral gavage on an HCT116 xenograft. Drugs were administrated at indicated days. **f,** an A549 xenograft model to compare the tumor suppression of auranofin-pantethine combo and cisplatin monotreatment. Drug administration was conducted two times per week with indicated concentrations. For **e-f**, data were shown as mean ± s. e. m. HCT116 tumors: Vehicle, n = 8 mice; pantethine, n = 7 mice; AF, n = 6 mice; AF + pantethine, n = 8 mice. A549 tumors: Vehicle, n = 6 mice; cisplatin, n = 6 mice; AF + pantethine, n = 4 mice.

### Pantethine potently synergizes auranofin both *in vitro* and *in vivo*

Pantethine, composed of pantothenate and cysteamine moieties (also the molecular fragments of CoA), is a dietary supplement for pantothenate-fueled CoA biosynthesis. Noteworthily, it can be orally administered at doses up to 1.2 g per day^53^, of which the good safety renders it a promising candidate as a clinical adjuvant for auranofin. Like TGTA, we found a dose-dependent auranofin-boosting manner of pantethine (Fig. 4b) along with an increase of cellular gold (Fig. 4c). To further verify the involvement of ligand exchange mechanism, we compared several structure-related molecules such as pantethine, pantetheine, cysteamine, and pantothenate for their abilities in synergizing auranofin. All these small molecules are related to the pantothenate metabolism^54^. For the lack of thiol group, pantothenate failed to boost auranofin (Fig. 4d). Pantetheine and its metabolic product cysteamine conferred a similar synergistic effect, and combo using cysteamine with pantothenate did not further enhance the effect (Fig. 4d). These results rule out the engagement of pantothenate metabolism in the synergistic effects, reinforcing a thiol-to-thiol exchange mechanism. Encouraged by the results above, we next performed the effects of the “auranofin + pantethine” combo in animal xenografts. Given that pantethine can be taken orally like auranofin, oral gavage was conducted in an HCT116 xenograft. As the results show (Fig. 4e), while single treatment by auranofin or pantethine did not show any tumor growth inhibition, the combo group led to significant inhibition, similar to the activity of TGTA with intraperitoneal injection. Likewise, the combo of auranofin-pantethine did not affect the body weight and organs of the treated mice (Extended Data Fig. 5a, b). Moreover, this orally available combo allowed the use of auranofin at 0.9 mg/kg, equivalent to the dosage permitted for treating RA in humans^55,56^, to significantly suppress tumor growth in an A549 xenograft. The activities of this combo are comparable to an intraperitoneal treatment of cisplatin at 4 mg/kg (Fig. 4f). No loss of body weight was found in this model as well (Extended Data Fig. 5a).

To better understand the influence of pantethine on gold ions in circulation, we monitored the plasma gold using auranofin-treated Sprague-Dawley rats. At initial, a single oral dose of auranofin (10 mg/kg) with or without pantethine (200 mg/kg) was administrated in each group. As the curves show, the gold content in the combo group tends to be slower than that of the auranofin alone group, albeit not significant (Extended Data Fig. 6). Since the gold ions can be maintained in blood for a long term (plasma half-lives of auranofin gold of 1.8 days in the rat, 19.5 days in the dog and 17 days in human) ^66–68^, we gave a second administration of pantethine (200 mg/kg) to the combo group after 24 hours. Significantly, the pantethine reduced the plasma gold levels, indicating a direct alteration in the albumin binding.

### DyThex drives auranofin to induce a cuproptosis-like cell death type

Besides the pharmacokinetics, next, we sought to investigate whether the DyThex nature of gold contributes to its intracellular mode of action. In the second-round ligand screening, lipoic acid (LA), the organosulfur derivative of octanoic acid, was found less effective in potentiating auranofin. Nonetheless, LA is endogenous and essential for metabolic regulation of the tricarboxylic acid (TCA) cycle via protein lipoylation^60^. It was recently identified that internalized copper ions can target those lipoylated proteins and induce a new type of cell death, coined cuproptosis^61^. Because of the high affinity of gold with thiols, it is possible for cellular gold(I) ions to dynamically interact with those disulfide-containing lipoylated proteins. Initially, we incubated auranofin with the intracellular form of lipoic acid (dihydrolipoic acid) ^62^, observing hybrid gold-lipoic acid adducts by HRMS (Fig. 5a). Next, we looked at the molecular markers identified in the cuproptosis study. FDX1 and LIAS are Fe-S cluster proteins involved in cuproptosis, showing a copper-dependent loss of protein levels. Then we observed a decline of FDX1 and LIAS levels in HCT116 cells upon auranofin treatment (Fig. 5b). The aggregate formation of DLAT protein, one of the lipoylated proteins, is another marker of cuproptosis^61^. As the blots show, auranofin can induce the aggregation of DLAT (Fig. 5c). In sum, the results suggested a cuproptosis-like mechanism induced by auranofin via gold-thiol exchanges.

**Fig. 5.**
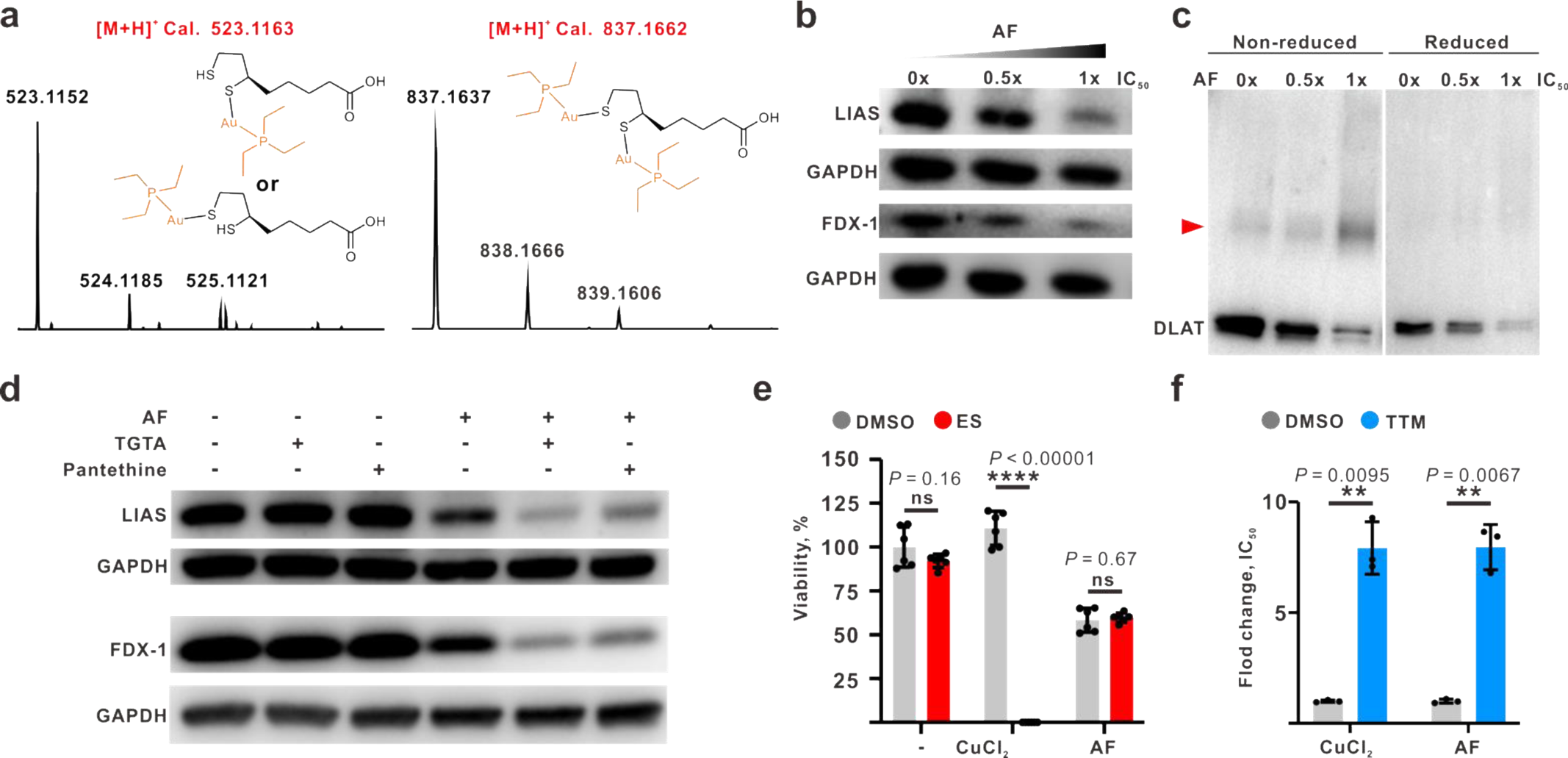
Auranofin induces cuproptosis-like cell death by thiol exchange. **a,** mass spectrometry identification of the products after thiol exchange in the incubation of auranofin and dihydrolipoic acid. **b,** analysis of the cuproptosis markers (LIAS and FDX1) in HCT116 cells under auranofin treatments. **c,** DLAT aggregation measurement by reduced and non-reduced blots under indicated treatments. the band of aggregated DLAT proteins was marked by a red arrow. **d,** the molecular markers of cuproptosis under auranofin mono or combined treatments as indicated. For **b-d,** each blot is representative of three biologically independent experiments. **e,** cell viability assay comparing the boost effect of elesclomol (ES) towards copper chloride and auranofin on HCT116. **f,** Effects of tetrathiomolybdate on the cytotoxicity of copper chloride and auranofin treatments on HCT116. The fold change of IC_50_ to that of the control group was shown. For **e-f**, data were presented as mean ± s. d. from 3 biologically independent replicates. Significance was calculated by unpaired, two-tailed *t*-test.

Compared to auranofin mono-treatment, combos using either TGTA or pantethine further suppressed the protein level of LIAS and FDX1 (Fig. 5d), in line with the aforementioned ligand-enhanced cytotoxicity. Of interest, it has been reported that copper-induced cell death is regulated by its ionophore^61^, or in other words, ligand^63^. Elesclomol (ES) dramatically promotes copper toxicity while tetrathiomolybdate (TTM) blocks^61^. By comparing copper chloride and auranofin, we found ES prefers to enhance copper (Fig. 5e). In contrast, the thiolate-containing TTM can detoxify both copper and gold (Fig. 5f), rendering a way to withdraw the gold cytotoxicity.

## Discussion

The daunting range of drug resistance in tumors continuously demands new drugs with orthogonal targeting to the current ones^64^. A highly bioactive metal center coupled with easy-to-release ligands makes metallodrug an efficient multitargeting agent, either used alone or as an adjuvant to other therapies^11^. However, the same ligand lability also results in a sophisticated behavior before they enter the tumor cells^12^, exemplified by the rapid thiol exchange with albumin affecting the tumor uptake of auranofin. In this work, we put forward a strategy to recover serum-inhibited auranofin simply by adding nontoxic thiol ligands, indicating the albumin-auranofin interaction is in a modulable dynamic equilibrium. Specifically, we confirmed both the TGTA ligand of auranofin and other biocompatible small-molecule thiols can boost the antitumor activities of auranofin in multiple tumor models. Of note, the orally administrated “auranofin + pantethine” combo produced an identical therapeutic outcome to cisplatin at 4 mg/kg in a mouse NSCLC model with a dosage of auranofin (0.9 mg/kg) equivalent to its approved use in rheumatoid arthritis. Together, our work illustrates that fast and dynamic ligand exchange can be utilized to realize an *in vivo* modulation on a metallodrug.

Two facets of our work may enlighten future studies in this field. Firstly, we illustrated how ligand-like small-molecule drugs can directly affect metallodrugs in a treatment regime. That is, the synergy between drugs may not only come from the target or pathway levels but also a direct ligand exchange in a rapid way *in vivo*. Ligand-mediated effects on auranofin in our work are determined by the chemical features of ligands, e.g., hydrophobicity. Since thiol is one of the most common groups in the bioactive molecules^65^, a pool of ligands with diverse chemical properties can be envisioned to boost or halt the gold’s toxicity to different degrees. That means an adjustable anticancer behavior even after the auranofin administration. Secondly, hydrophilic thiols but preferred by some cell types may also potentiate auranofin in a much more selective manner. Encouraged by our discovery that auranofin plus CoA only works in the Jurkat T cell model, comparing the gene expression or dependency levels of Jurkat with other cell lines using well-established databases like DepMap^18,57^ could potentially unleash some hints for such a cell-specific activity of CoA. Subsequent studies to understand the cancer-enriched thiols may be fruitful as well. On the other hand, attaching thiol groups to already-in-use tumor-targeting agents such as small molecules, peptides, antibodies, and nanoparticles seems a more feasible way for developing highly selective and effective anticancer combos with auranofin.

More than the case of repurposing auranofin, our methods may also be applied to other metallodrugs since many of them possess labile ligands and the subsequent unpredictable manners in the serum. For the developing drug candidates, screening relevant ligands directly with the initial metal complex obviates the need for a labor-intensive synthesis of structural derivatives. For those already used metallodrugs, knowledge of the modulable nature highlighted in this work may be instructive to clinicians who are considering delicate adjustments of the activity and side toxicity using drug combos. Thus, we believe this work is of importance to the future development of metallodrugs either in mono- or combined therapies.

## Methods

### Ethics statement

The research performed in the present study complies with all ethical regulations. The animal procedures were approved by the Research Ethics Committee of Sun Yat-Sen University (SYSU-IACUC-2020-0826) and conducted following the Guide for the Care and Use of Laboratory. All animals were purchased from the Experimental Animal Center of Sun Yat-Sen University (Guangzhou, China) and Nanjing Biomedical Research Institute of Nanjing University (Nanjing, China). All mice and rats were housed under specific-pathogen-free conditions and maintained in single-sex cages at 23 ± 3 °C and 40–70% humidity under a 12/12-h light/dark cycle. Inclusion criteria were: male or female, appropriate age and weight (15–30 g). Exclusion criteria were: tumor size must not exceed 15 mm (volume 2,000 mm^3^) in any direction in an adult mouse, tumor mass should not proceed to the point where it significantly interferes with normal bodily functions or causes pain or distress owing to its location, persistent self-induced trauma, rapid or progressive weight loss of more than 25%, for seven days. In none of the experiments were these approved ethical limits exceeded.

### Cell lines and cell culture

Human cancer cell lines including A549 (lung carcinoma cells), MDA-MB-231 (triple-negative breast cancer cells), HepG2 (hepatoma carcinoma cells), A375 (melanoma cells), PANC-1 (pancreatic carcinoma cells) and SW480 (colon adenocarcinoma cells) were from the American Type Culture Collection (ATCC). PC9 (lung carcinoma cells) were from the European Collection of Authenticated Cell Cultures (ECACC, London, UK). HCT116 (colon adenocarcinoma cells) were from Procell company (Wuhan, China). Jurkat (human T lymphoblastic leukemia cells) were generously gifted by Guohui Wan at Sun Yat-Sen University. U87 (glioblastoma cells) were generously provided by Min Feng group at Sun Yat-Sen University. PC9, A549, HCT116, and Jurkat were cultured using RPMI 1640 (Corning). HepG2, A375, PANC-1, SW480, MDA-MB-231, and U87 cells were cultured using DMEM (Thermo Fisher Scientific). All media were supplemented with 10% fetal bovine serum (Hyclone) and 1% penicillin-streptomycin G (Thermo Fisher Scientific).

### Chemicals

Auranofin, 1-thio-β-D-glucose tetraacetate (TGTA), D-pantethine, R-pantetheine, cysteamine, calcium pantothenate, ammonium tetrathiomolybdate and bovine serum albumin (BSA) were purchased from Sigma-Aldrich. Disulfiram, β-D-glucose pentaacetate, coenzyme A, lipoic acid, dihydrolipoic acid, oltipraz, pyrithioxin dihydrochloride, and cisplatin were purchased from MedChemExpress. 1-thio-β-D-Glucose (sodium salt) was purchased from Cayman. Elesclomol was purchased from TargetMol. Captopril was purchased from Selleck. Cremophor EL was purchased from Millipore. All chemicals were stored as stock solutions in DMSO or ultrapure water.

### Mouse studies

All animal studies and procedures were conducted according to a protocol approved by the Research Ethics Committee of Sun Yat-Sen University. Mice were blindly randomized into different groups for treatment studies. Tumor volume was calculated with the equation V = (length × width^2^)/2 and mouse body weight was also monitored during the study. For effects on PDX models, both the lung cancer PDX sample (ID: TM00192) and breast cancer PDX sample (ID: TM01273F707) are from the Jackson Laboratory, and we followed the previous protocol for PDX transplantation^58^. Lung cancer PDXs were transplanted by inserting 2 mm^3^ into the dorsal flank of 4-6-week-old male NOD/SCID mice; breast cancer PDXs were transplanted by inserting 2 mm^3^ into the dorsal flank of 4-6-week-old female NOD/SCID mice. When the volume of cancer PDX was approximately 40 mm^3^, the mice were randomized and then treated intraperitoneally with 100 μL of either vehicle or 40 mg/kg TGTA (in corn oil) or 3 mg/kg AF (in a formulation of 15% Cremophor EL, 82.5% PBS and 2.5% DMSO) alone or combination treatment for 6 times per week.

For the establishment of HCT116 and A549 xenografts, 4-6-week-old male BALB/c nu/nu athymic mice with a body weight of 18–23 g were used. For evaluation of the effects of ‘AF+TGTA’ on human colon cell line xenografts, HCT116 cells were injected (2 × 10^6^ cells transplanted subcutaneously) to grow tumors. When tumor volume had reached around 40 mm^3^, mice were randomized into four treatment groups. The mice were then treated intraperitoneally with 100 μL of either vehicle or 40 mg/kg TGTA or 3 mg/kg AF alone or combination treatment 6 times per week. One week was considered as one cycle.

For evaluation of the effects of ‘AF+DPT’ on human colon cell line xenografts, HCT116 cells were injected (2 × 10^6^ cells transplanted subcutaneously) to grow tumors. When tumor volume had reached around 40 mm^3^, mice were randomized into four treatment groups. The mice were then treated by oral gavage with 100 μL of either vehicle or 200 mg/kg D-pantethine (in saline) or 3 mg/kg AF alone or combination treatment. For evaluation of the effects of ‘AF+DPT’ on human lung cell line xenografts, A549 cells were injected (4 × 10^6^ cells transplanted subcutaneously) to grow tumors. When tumor volume had reached around 40 mm^3^, mice were randomized into three treatment groups. Mice were given 0.9 mg/kg AF and 200 mg/kg D-pantethine for combination treatment by oral gavage 2 times per week. The positive group was treated intraperitoneally with 100 μL of 4 mg/kg cisplatin (in saline) 2 times per week. One week was considered as one cycle.

### Plasma gold studies in rats

Ten male Sprague-Dawley rats were equally divided into two groups. After fasting for 24 hours, the rats were given a single oral dose of auranofin (10 mg/kg) with or without pantethine (200 mg/kg). After 24 hours, the rats were treated orally with either saline or pantethine (200 mg/kg) again. The rats were placed in cages and 0.5 mL of heparinized blood was collected from each rat at 1.5, 6, 10, 24, 30, and 34 hours after administration of the drug. Approximately 0.5 mL of each blood specimen was transferred to a clean tube and centrifuged at 400 xg for 20 minutes. Plasma from each specimen was transferred to a clean tube with a dropping pipette. The collected plasma was digested with mixed acid (HNO_3_:HClO_4_ = 3:1) at 95 °C overnight. The same volume of ultrapure water was transferred into a separate EP tube for the background subtraction. Then, the concentration of Au in the plasma was examined with inductively coupled plasma mass spectrometry (ICP-MS) analysis.

### Cell viability and growth assays

Cells were seeded in 96-well plates at 6,000 cells per well and cultured for 24 hours before being subjected to various treatments. For the in-house ligand library screening, PC9 cells were used. Cells were treated with various ligands (50 µM) in the absence or presence of 5 μM AF in RPMI 1640 media supplemented with 30% fetal bovine serum. For the small-scale cell line profiling of pantethine, pantetheine, CoA, lipoic acid, and thioglucose, all cell lines were cultured in the human plasma-like medium (Gibco^TM^) supplemented with 30% fetal bovine serum. After a total of 24 h treatment with indicated agents, cell viability was assessed using 20 μL of MTT (3-(4,5-dimethyl-2-thiazolyl)-2,5-diphenyl tetrazolium bromide) solutions and measured on a microplate spectrophotometer at a wavelength of 490 nm.

### High-performance liquid chromatograms (HPLC)

High-performance liquid chromatograms were conducted with a reversed-phase C_18_ column at 214 nm to monitor the gold-thiol adducts^59^. The mobile phase consisted of a 65:35 solution of methanol-water with a flow rate of 1.0 ml/min. The column (length, 15 cm; inner diameter, 4.6 mm) was a Cosmosil model commercially packed with octadecylsilane reversed-phase material (code 38019-81). Experiments were performed at constant temperature (30 °C). Portions of 0.25 mol equivalent of a solution of auranofin (100 mM in 10 µL MeOH) were added to 1.0 mL of 4 mM solution of BSA and then reacted at 37 °C for 60 min in a shaker moving. The sample was divided into two aliquots of 250 µL each, one with 5.0 eq. TGTA (50 mM in MeOH) and one with an equal volume of methanol and the reaction was carried out for 20 minutes. Then all samples were precipitated by adding 4 times the amount of acetone (1.0 mL) and centrifuged at high speed for 10 minutes. The supernatant was filtered and then loaded into HPLC. All solutions were filtered through 0.22 µm membrane filters before use.

### Determination of Cellular TrxR activity

The cellular TrxR experiment was performed by using the Thioredoxin Reductase Activity Assay Kit (Solarbio) ^69^. HCT116 cells were inoculated into a 6-well plate at the density of 2×10^5^ cells per well and incubated for 36 h before being subjected to various treatments. After 1 h of incubation, the cells were washed three times with ice-cold PBS and then were added with 100 μL of ice-cold lysis buffer (50 mM phosphate buffer, pH 7.4, 1 mM EDTA, 0.1% Triton-X100). The protein concentrations were determined by BCA assay (Yeasen). Cellular TrxR activity was measured by a microplate Reader (OD 412 nm) according to the manufacturer’s instructions.

### ICP-MS experiments

For quantitative analysis of cellular uptake, U87 cells were seeded in 4 cm dishes at a density of 1×10^5^ cells/ml (4.0 ml) and allowed to attach for 24 h. The cells were then treated with auranofin, disulfiram, and captopril (1:190:260) (3 µM) for 10 min at 37 °C in a CO_2_ incubator. At the end of the incubation, cells were collected and washed three times with ice-cold PBS. The collected cells were digested with mixed acid (HNO_3_:HClO_4_ = 3:1) at 95 °C overnight. The same volume of ultrapure water was transferred into a separate EP tube for the background subtraction, at least in duplicate per experiment. Then, the concentration of Au in the cells was examined with inductively coupled plasma mass spectrometry (ICP-MS) analysis. Values were normalized against protein content.

### Antibodies and western blotting

The following antibodies were used: polyclonal rabbit anti-LIAS (1:1,000; catalog no. 11577-1-AP; Proteintech); monoclonal rabbit anti-FDX1 (1:1,000; catalog no. ab108257; Abcam); monoclonal mouse anti-DLAT (1:1,000; catalog no. 12362S; Cell signaling Technology); monoclonal mouse HRP-conjugated GAPDH (1:10,000; catalog no. HRP-60004; Proteintech); polyclonal HRP-conjugated affinipure goat anti-mouse IgG(H+L) (1:10,000; catalog no. SA00001-1; Proteintech) and polyclonal HRP-conjugated affinipure goat anti-rabbit IgG(H+L) (1:10,000; catalog no. SA00001-2; Proteintech). HCT116 cells were inoculated into 4 cm tissue culture plates and grown to 75% confluence before being subjected to various treatments. The cells were washed three times with ice-cold PBS and then were added with 100 μL of ice-cold lysis buffer supplemented by protease and phosphatase inhibitors (Sigma-Aldrich). Lysates were centrifuged at 10,000 xg at 4 °C for 12 min, and the protein concentration of the supernatant was determined using the BCA Protein Assay. 40 μg of protein per sample was mixed with sample buffer and denatured at 95 °C for 5 min. Samples were separated by SDS–PAGE and transferred to a nitrocellulose membrane. Membranes were blocked with 5% skim milk (Biofroxx) for 1 h at room temperature and then incubated with a 1:1,000 dilution of antibody at 4 °C overnight. Secondary antibodies were then incubated with the membrane for 1 h at room temperature, and the membrane was imaged using the Tanon Imaging System.

### High-resolution mass spectrometry (HRMS)

HRMS solutions were prepared and injected without further dilution. Stock solution of AF:disulfiram:captopril (1:1:2) (2 μM) was prepared in analytical grade methanol. The HRMS solutions were prepared in methanol (1.0 mL) with AF:dihydrolipoic acid (DHLA) (5:1) (0.8 µM). The HRMS solutions of albumin-Au-PEt3 adduct were prepared in ammonium acetate (2 mM) with AF:BSA (8:1) (6 µM).

---

**Extended Data Fig. 1.**
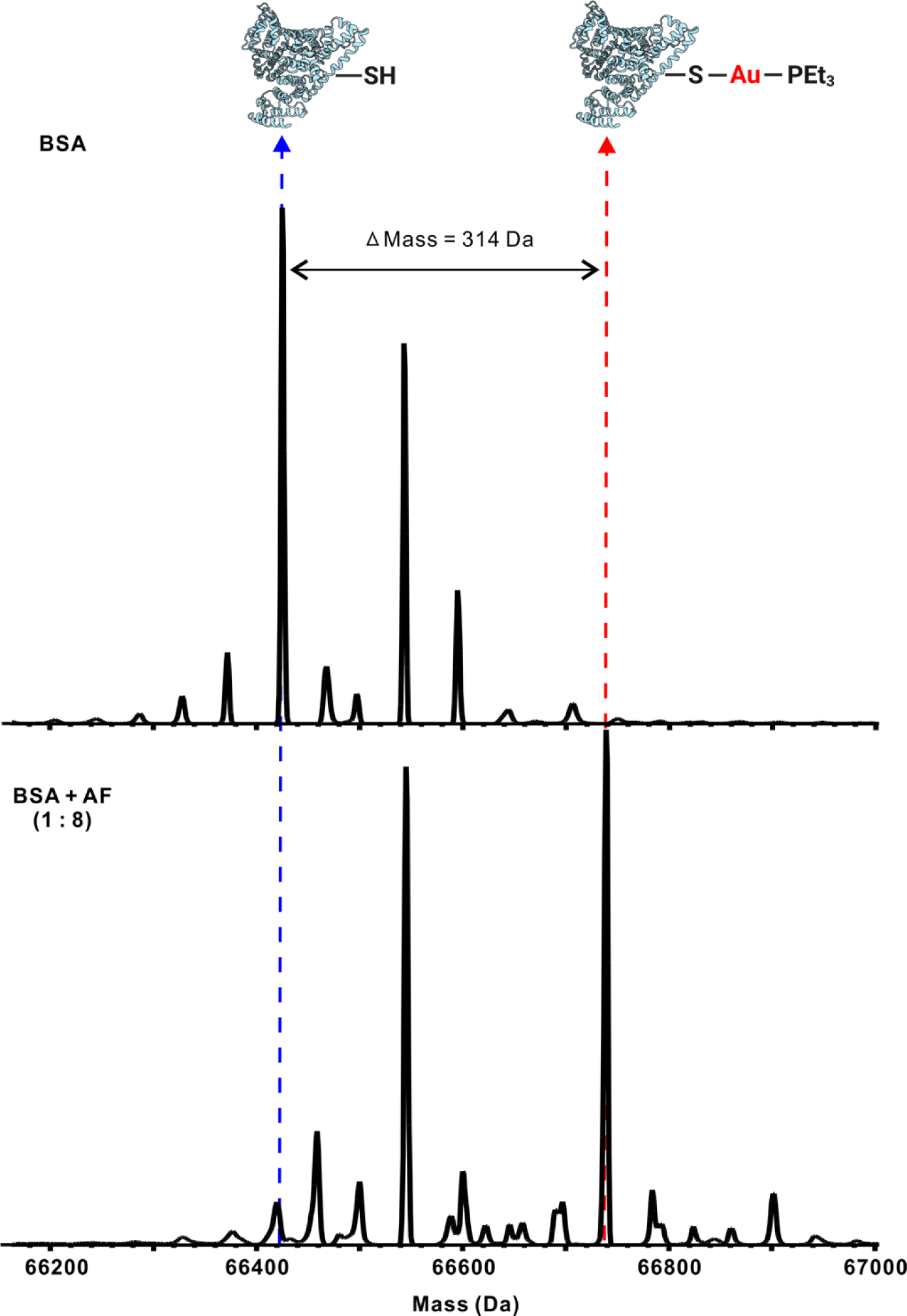
Analysis of auranofin-albumin complex. MALDI-TOF analysis of the product of coincubation of auranofin and albumin. The gold-albumin adduct was marked by a red arrow while the peak of albumin was marked by a blue arrow. A shift of 314 Da represents the addition of a gold-phosphine group that binds to the Cys residue of albumin by thiol exchange.

**Extended Data Fig. 2.**
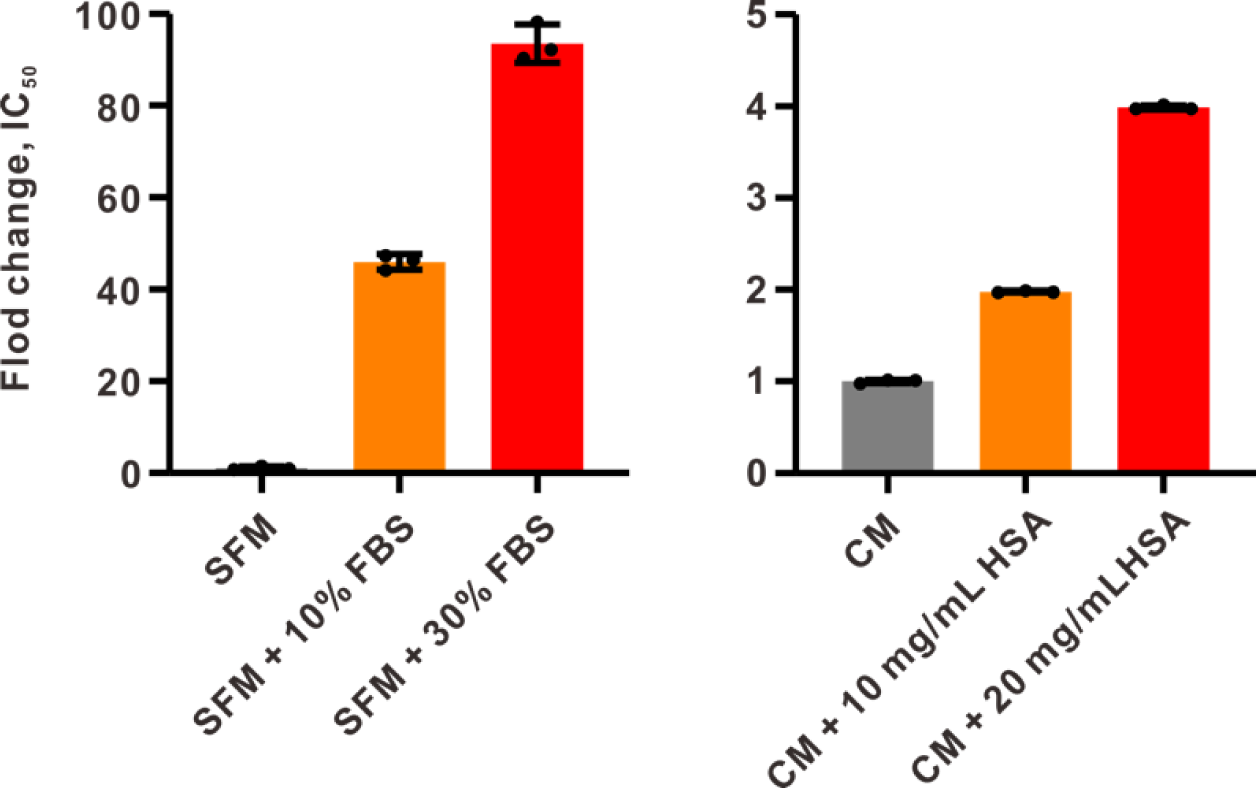
Auranofin cytotoxicity blocked by serum and albumin. Fold changes of the IC_50_ value of auranofin towards HCT116 cells upon indicated FBS or HSA conditions were shown. FBS, fetal bovine serum; HSA, human serum albumin; SFM, serum-free medium (RPMI 1640); CM, culture medium (RPMI 1640 + 10% FBS). Data are shown as mean ± s. d. of three independent experiments.

**Extended Data Fig. 3.**
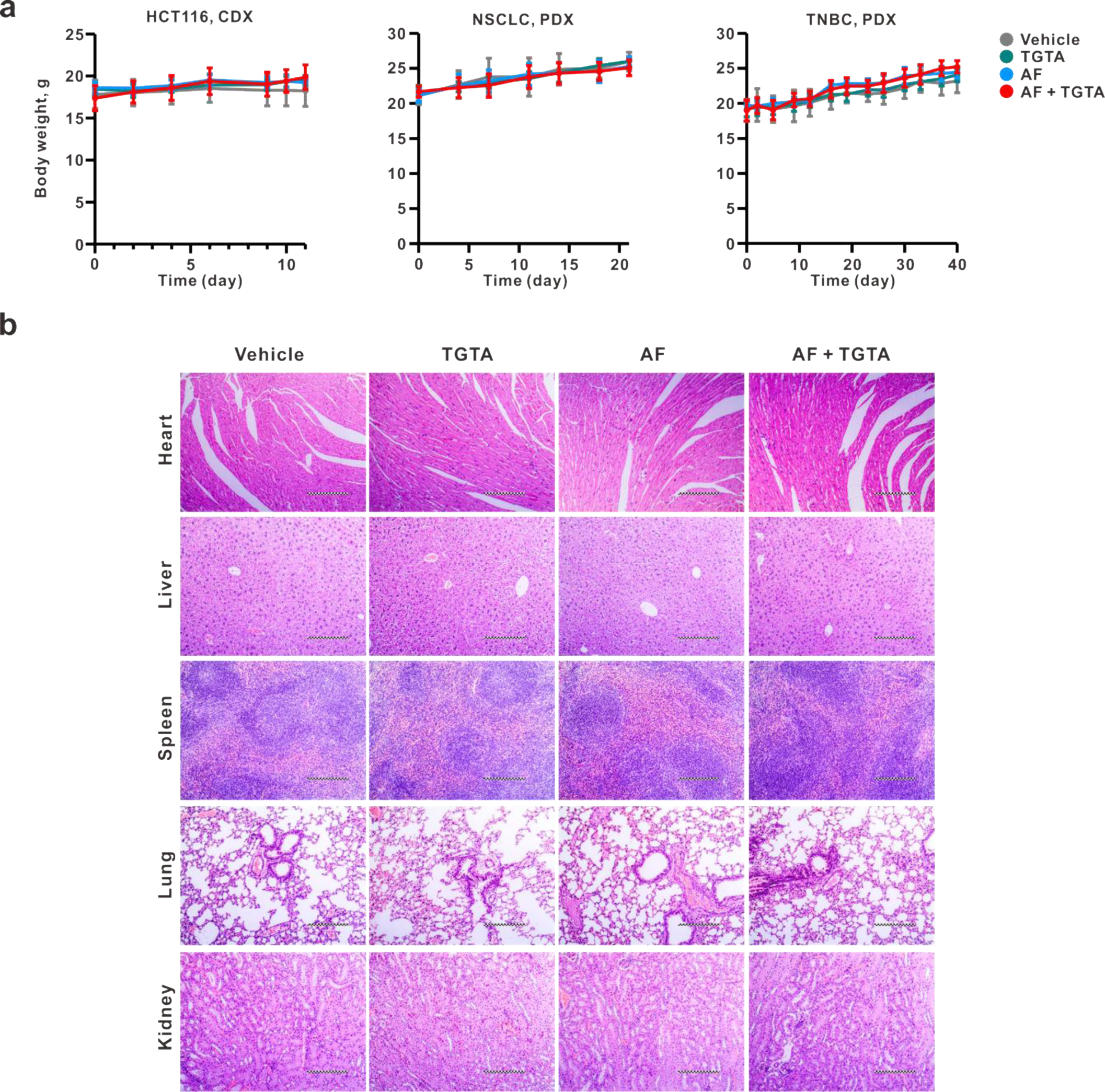
Impacts of auranofin-TGTA combo on the mouse physiology. **a,** monitoring the mouse body weight of the three models, HCT116 CDX, NSCLC PDX, and TNBC PDX. **b,** pathological analysis of hearts, kidneys, livers, lungs and spleens harvested from the NSCLC PDX models by hematoxylin-eosin staining. Each picture is a representative of three biologically independent replicates.

**Extended Data Fig. 4.**
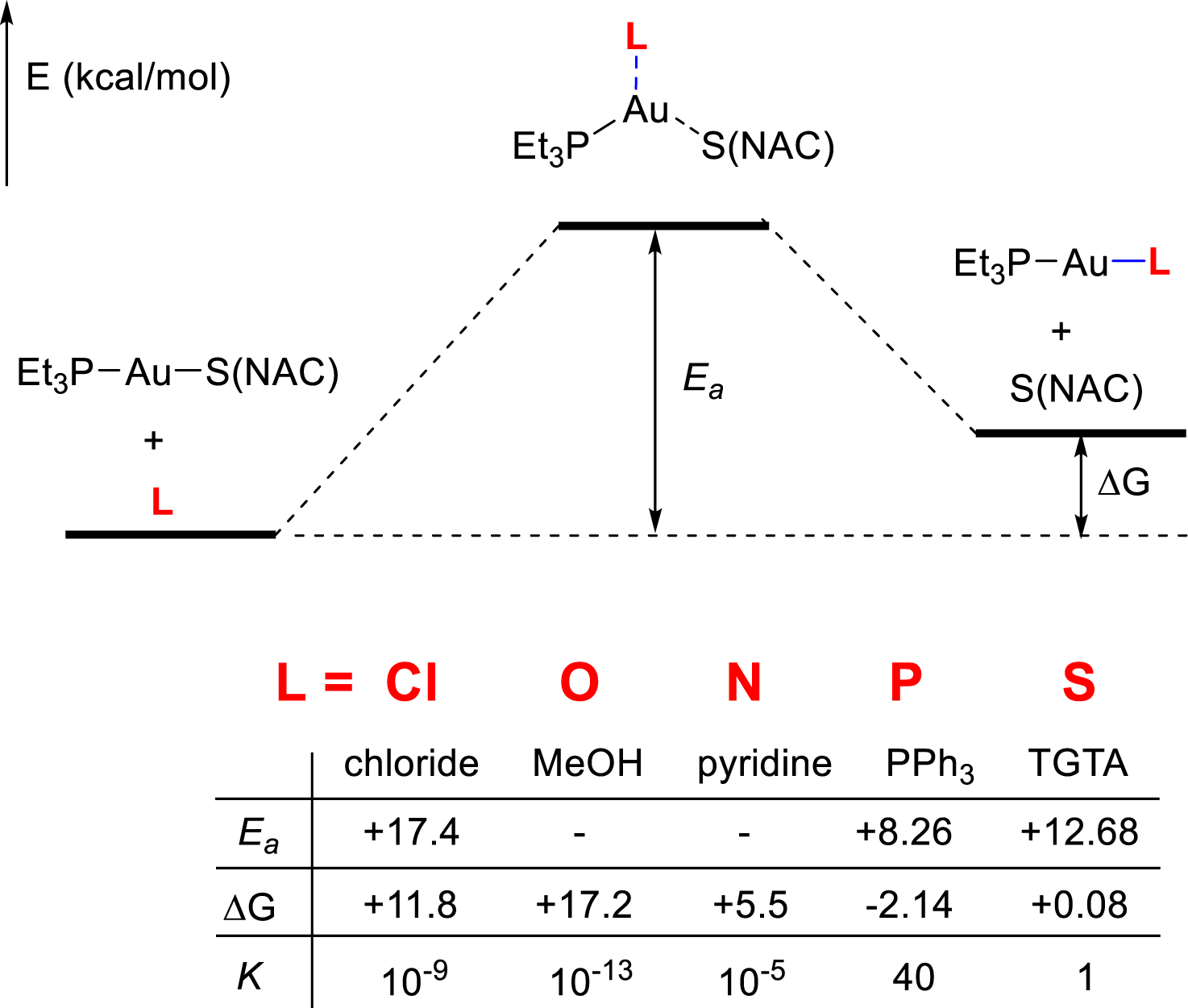
Computational evaluation (DFT) of the exchange reactions with albumin-bound gold by different ligand types. *E_a_*, activation energy; ΔG, Gibbs free energy; *K*, chemical equilibrium constant calculated from ΔG. Note: the gold compounds were optimized by the CAM-B3LYP functional, with basis set of SDD for Au and 6-31G* for others. The single point energies were calculated at the same level with SDD basis set for Au and 6-311++G** for others. Water was set as the solvent model based on the SMD model.

**Extended Data Fig. 5.**
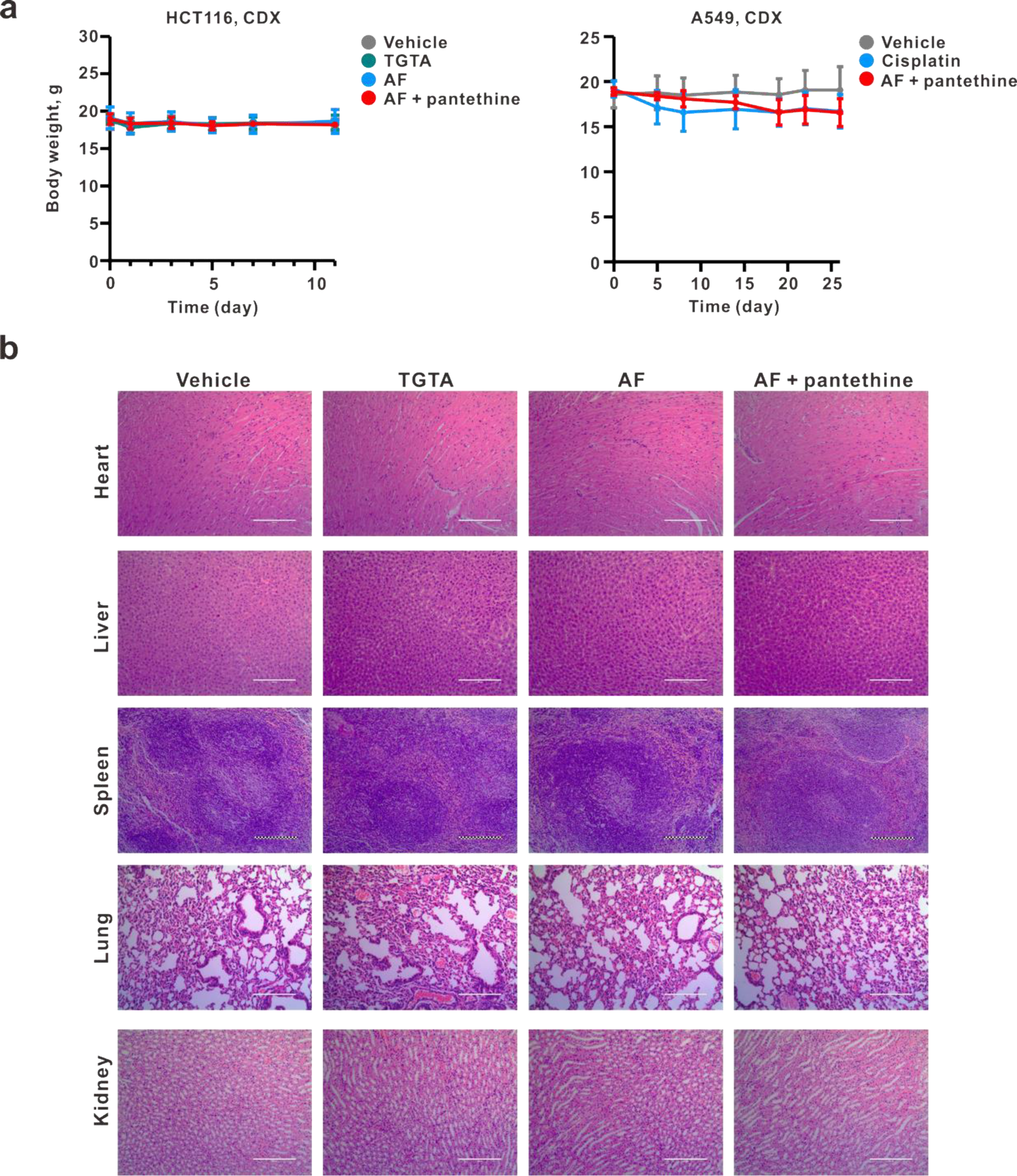
Impacts of auranofin-pantethine combo on the mouse physiology. **a,** monitoring the mouse body weight of the two models, HCT116 CDX, and A549 PDX. **b,** pathological analysis of hearts, kidneys, livers, lungs and spleens harvested from the HCT116 xenograft models by hematoxylin-eosin staining, under the indicated treatments. Each picture is a representative of three biologically independent replicates.

**Extended Data Fig. 6.**
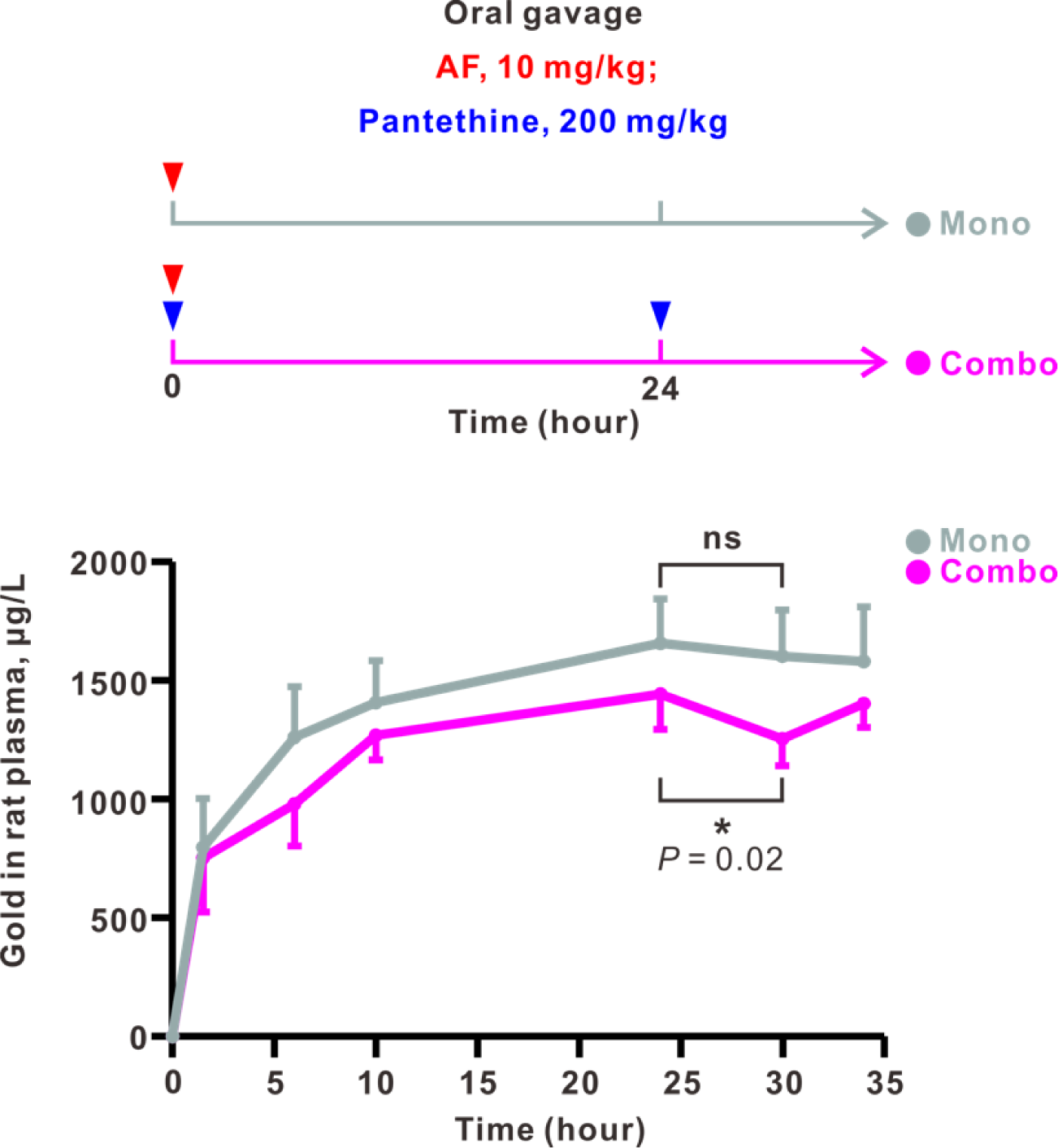
Monitoring the plasma gold in a rat model under indicated drug administrations. Curves represent the plasma gold concentration measured under the treatment of auranofin alone (Mono) and auranofin-pantethine group (Combo), respectively. The upper part shows the drug administration conditions for each group. The timepoints of drug administration were marked by arrows with indicated colors for each drug. For each administration, auranofin (AF) was used at 10 mg/kg and pantethine at 200 mg/kg. The significance indicated was calculated by paired, two-tailed *t*-test.

## Acknowledgements

This work was finically supported by the National Natural Science Foundation of China (nos. 22122706, 22377154 and 32100058), Guangdong Science and Technology Department (no. 2019QN01C125), Guangdong Basic and Applied Basic Research Foundation (nos. 2021A1515012347, 2021A1515011168, and 2020A1515110508), and Guangdong Provincial Key Laboratory of Construction Foundation (no. 2023B1212060022).

## Author Information

These authors contributed equally: Yuan Wang, Bei Cao and Qianqian Wang.

## Contributions

T.Z. conceived and designed the study. Y.W., B.C., Q.W., X.F., J.W., X.X. and T.Z. designed and performed the experiments. X.X., Y.W., and T.Z. contributed to the writing and editing of the manuscript and all authors approved the final edition of the manuscript. Supervision was provided by J.W., X.X., and T.Z.

## Ethics declarations

### Data availability

Source data are provided in this paper. All other data supporting the findings of this study are available from the corresponding author upon reasonable request. Requests for experimental agents and materials will be reviewed promptly by the corresponding author.

## References

1 Rottenberg, S., Disler, C. & Perego, P. The rediscovery of platinum-based cancer therapy. Nat. Rev. Cancer 21, 37–50 (2021).

2 Forde, P. M., Spicer, J., Lu, S., Provencio, M., Mitsudomi, T., Awad, M. M., Felip, E., Broderick, S. R., Brahmer, J. R., Swanson, S. J., Kerr, K., Wang, C. L., Ciuleanu, T. E., Saylors, G. B., Tanaka, F., Ito, H., Chen, K. N., Liberman, M., Vokes, E. E., Taube, J. M., Dorange, C., Cai, J. L., Fiore, J., Jarkowski, A., Balli, D., Sausen, M., Pandya, D., Calvet, C. Y. & Girard, N. Neoadjuvant Nivolumab plus Chemotherapy in Resectable Lung Cancer. N. Engl. J. Med. 386, 1973–1985 (2022).

3 Wang, Z. X., Cui, C. X., Yao, J., Zhang, Y. Q., Li, M. X., Feng, J. F., Yang, S. J., Fan, Y., Shi, J. H., Zhang, X. Z., Shen, L., Shu, Y. Q., Wang, C. L., Dai, T. Y., Mao, T., Chen, L., Guo, Z. Q., Liu, B., Pan, H. M., Cang, S. D., Jiang, Y., Wang, J. Y., Ye, M., Chen, Z. D., Jiang, D., Lin, Q., Ren, W., Wang, J. S., Wu, L., Xu, Y., Miao, Z. H., Sun, M. L., Xie, C. H., Liu, Y., Wang, Q. F., Zhao, L. N., Li, Q., Huang, C. H., Jiang, K., Yang, K. Y., Li, D. J., Liu, Y. P., Zhu, Z. T., Chen, R. X., Jia, L. Q., Li, W., Liao, W. J., Liu, H. X., Ma, D. Y., Ma, J., Qin, Y. R., Shi, Z. H., Wei, Q. C., Xiao, K., Zhang, Y., Zhang, Y., Chen, X., Dai, G. H., He, J. X., Li, J. H., Li, G. H., Liu, Y., Liu, Z. H., Yuan, X. L., Zhang, J. P., Fu, Z. C., He, Y. F., Ju, F., Liu, Z., Tang, P., Wang, T. J., Wang, W. B., Zhang, J., Luo, X. M., Tang, X. W., May, R., Feng, H., Yao, S., Keegan, P., Xu, R. H. & Wang, F. Toripalimab plus chemotherapy in treatment-naive, advanced esophageal squamous cell carcinoma (JUPITER-06): A multi-center phase 3 trial. Cancer Cell 40, 277–288 (2022).

4 Liu, P., Chen, J. Z., Zhao, L. W., Hollebecque, A., Kepp, O., Zitvogel, L. & Kroemer, G. PD-1 blockade synergizes with oxaliplatin-based, but not cisplatin-based, chemotherapy of gastric cancer. Oncoimmunology 11, 2093518 (2022).

5 Pfirschke, C., Engblom, C., Rickelt, S., Cortez-Retamozo, V., Garris, C., Pucci, F., Yamazaki, T., Poirier-Colame, V., Newton, A., Redouane, Y., Lin, Y. J., Wojtkiewicz, G., Iwamoto, Y., Mino-Kenudson, M., Huynh, T. G., Hynes, R. O., Freeman, G. J., Kroemer, G., Zitvogel, L., Weissleder, R. & Pittet, M. J. Immunogenic Chemotherapy Sensitizes Tumors to Checkpoint Blockade Therapy. Immunity 44, 343–354 (2016).

6 Petroni, G., Kroemer, G. & Galluzzi, L. Immunogenic Therapies Drive CAR T Cells towards Superior Efficacy. Trends Cancer 7, 179–181 (2021).

7. Bertini, I. G., Harry B.; Lippard, Stephen J.; Valentine, Joan Selverstone. Bioinorganic Chemistry (University Science Books, 1994).

8 Nagai, N., Okuda, R., Kinoshita, M. & Ogata, H. Decomposition kinetics of cisplatin in human biological fluids. J. Pharm. Pharmacol. 48, 918–924 (1996).

9 Wang, D. & Lippard, S. J. Cellular processing of platinum anticancer drugs. Nat. Rev. Drug Discovery 4, 307–320 (2005).

10 Cini, M., Bradshaw, T. D. & Woodward, S. Using titanium complexes to defeat cancer: the view from the shoulders of titans. Chem. Soc. Rev. 46, 1040–1051 (2017).

11 Anthony, E. J., Bolitho, E. M., Bridgewater, H. E., Carter, O. W. L., Donnelly, J. M., Imberti, C., Lant, E. C., Lermyte, F., Needham, R. J., Palau, M., Sadler, P. J., Shi, H. Y., Wang, F. X., Zhang, W. Y. & Zhang, Z. J. Metallodrugs are unique: opportunities and challenges of discovery and development. Chem. Sci. 11, 12888–12917 (2020).

12 Riccardi, L., Genna, V. & De Vivo, M. Metal-ligand interactions in drug design. Nat. Rev. Chem. 2, 100–112 (2018).

13 Zou, T. T., Lok, C. N., Wan, P. K., Zhang, Z. F., Fung, S. K. & Che, C. M. Anticancer metal-N-heterocyclic carbene complexes of gold, platinum and palladium. Curr. Opin. Chem. Biol. 43, 30–36 (2018).

14 Jiang, J., Xiong, X. L. & Zou, T. T. Modulating the Chemical Reactivity of Gold Complexes in Living Systems: From Concept to Biomedical Applications. Acc. Chem. Res. 56, 1043–1056 (2023).

15 Corsello, S. M., Bittker, J. A., Liu, Z. H., Gould, J., McCarren, P., Hirschman, J. E., Johnston, S. E., Vrcic, A., Wong, B., Khan, M., Asiedu, J., Narayan, R., Mader, C. C., Subramanian, A. & Golub, T. R. The Drug Repurposing Hub: a next-generation drug library and information resource. Nat. Med. 23, 405–408 (2017).

16 Beijersbergen, R. L. Old drugs with new tricks. *Nat*. Cancer 1, 153–155 (2020).

17 Gyawali, B., Bouche, G., Crisp, N. & Andre, N. Challenges and opportunities for cancer clinical trials in low- and middle-income countries. *Nat*. Cancer 1, 142–145 (2020).

18 Corsello, S. M., Nagari, R. T., Spangler, R. D., Rossen, J., Kocak, M., Bryan, J. G., Humeidi, R., Peck, D., Wu, X., Tang, A. A., Wang, V. M., Bender, S. A., Lemire, E., Narayan, R., Montgomery, P., Ben-David, U., Garvie, C. W., Chen, Y., Rees, M. G., Lyons, N. J., McFarland, J. M., Wong, B. T., Wang, L., Dumont, N., O’Hearn, P. J., Stefan, E., Doench, J. G., Harrington, C. N., Greulich, H., Meyerson, M., Vazquez, F., Subramanian, A., Roth, J. A., Bittker, J. A., Boehm, J. S., Mader, C. C., Tsherniak, A. & Golub, T. R. Discovering the anti-cancer potential of non-oncology drugs by systematic viability profiling. *Nat*. Cancer 1, 235–248 (2020).

19 Yuan, S. F., Wang, R. M., Chan, J. F. W., Zhang, A. J. X., Cheng, T. F., Chik, K. K. H., Ye, Z. W., Wang, S. Y., Lee, A. C. Y., Jin, L. J., Li, H. Y., Jin, D. Y., Yuen, K. Y. & Sun, H. Z. Metallodrug ranitidine bismuth citrate suppresses SARS-CoV-2 replication and relieves virus-associated pneumonia in Syrian hamsters. Nat. Microbiol. 5, 1439–1448 (2020).

20 Hatem, E., Azzi, S., El Banna, N., He, T. T., Heneman-Masurel, A., Vernis, L., Baille, D., Masson, V., Dingli, F., Loew, D., Azzarone, B., Eid, P., Baldacci, G. & Huang, M. E. Auranofin/Vitamin C: A Novel Drug Combination Targeting Triple-Negative Breast Cancer. J. Natl Cancer Inst. 111, 597–608 (2019).

21. Lee, D., Xu, I. M. J., Chiu, D. K. C., Leibold, J., Tse, A. P. W., Bao, M. H. R., Yuen, V. W. H., Chan, C. Y. K., Lai, R. K. H., Chin, D. W. C., Chan, D. F. F., Cheung, T. T., Chok, S. H., Wong, C. M., Lowe, S. W., Ng, I. O. L. & Wong, C. C. L. Induction of Oxidative Stress Through Inhibition of Thioredoxin Reductase 1 Is an Effective Therapeutic Approach for Hepatocellular Carcinoma. Hepatology 69, 1768–1786 (2019).

22 Shen, S. Y., Shen, J., Luo, Z., Wang, F. D. & Min, J. X. Molecular mechanisms and clinical implications of the gold drug auranofin. Coord. Chem. Rev. 493, 215323 (2023).

23 Nobili, S., Mini, E., Landini, I., Gabbiani, C., Casini, A., Messori, L. Gold Compounds as Anticancer Agents: Chemistry, Cellular Pharmacology and Preclinical Studies. Med. Res. Rev. 30, 550–580 (2010).

24. Berners-Price, S. J., Mirabelli, C. K., Johnson, R. K., Mattern, M. R., McCabe, F. L., Faucette, L. F., Sung, C.-M., Mong, S.-M., Sadler, P. J., Crooke, S. T. In Vivo Antitumor Activity and in Vitro Cytotoxic Properties of Bis[1,2-bis(diphenylphosphino)ethane]gold(I) Chloride. Cancer Res. 46, 5486–5493 (1986).

25. Moreno-Alcántar, G., Picchetti, P., Casini, A. Gold Complexes in Anticancer Therapy: From New Design Principles to Particle-Based Delivery Systems. Angew. Chem. Int. Ed. 62, e202218000 (2023).

26 Milacic, V., Chen, D.; Ronconi, L., Landis-Piwowar, K. R., Fregona, D., Dou, Q. P. A Novel Anticancer Gold(III) Dithiocarbamate Compound Inhibits the Activity of a Purified 20S Proteasome and 26S Proteasome in Human Breast Cancer Cell Cultures and Xenografts. Cancer Res. 66, 10478–10486(2006).

27. Zhang, J.-J., Abu el Maaty, M. A., Hoffmeister, H., Schmidt, C., Muenzner, J. K., Schobert, R., Wölfl, S.; Ott, I. A Multitarget Gold(I) Complex Induces Cytotoxicity Related to Aneuploidy in HCT-116 Colorectal Carcinoma Cells. Angew. Chem. Int. Ed. 59, 16795–16800 (2020).

28 Ott, I. On the Medicinal Chemistry of Gold Complexes as Anticancer Drugs. Coord. Chem. Rev. 253, 1670–1681(2009).

29 Sadler, P. J., Sue, R. E. The Chemistry of Gold Drugs. Met.-Based Drugs 1, 107–144(1994).

30 Shaw III, C. F. Gold-Based Therapeutic Agents. Chem. Rev. 99, 2589–2600 (1999).

31 Cosottini, L., Massai, L., Ghini, V., Zineddu, S., Geri, A., Mannelli, M., Ciambellotti, S., Severi, M., Gamberi, T., Messori, L., Turano, P. Bioconjugation of the gold drug auranofin to human ferritin yields a potent cytotoxin. J. Drug. Deliv. Sci. Tec. 87, 104822–104831 (2023).

32 Hickey, J. L., Ruhayel, R. A., Barnard, P. J., Baker, M. V., Berners-Price, S. J. & Filipovska, A. Mitochondria-targeted chemotherapeutics: The rational design of gold(I) N-heterocyclic carbene complexes that are selectively toxic to cancer cells and target protein selenols in preference to thiols. J. Am. Chem. Soc. 130, 12570–12571 (2008).

33 Ralph, S. J., Nozuhur, S., Ra, A. L., Rodriguez-Enriquez, S. & Moreno-Sanchez, R. Repurposing drugs as pro-oxidant redox modifiers to eliminate cancer stem cells and improve the treatment of advanced stage cancers. Med. Res. Rev. 39, 2397–2426 (2019).

34 Gamberi, T., Chiappetta, G., Fiaschi, T., Modesti, A., Sorbi, F. & Magherini, F. Upgrade of an old drug: Auranofin in innovative cancer therapies to overcome drug resistance and to increase drug effectiveness. Med. Res. Rev. 42, 1111–1146 (2022).

35 Zhang, J. B., Simpson, C. M., Berner, J., Chong, H. B., Fang, J. F., Ordulu, Z., Weiss-Sadan, T., Possemato, A. P., Harry, S., Takahashi, M., Yang, T. Y., Richter, M., Patel, H., Smith, A. E., Carlin, A. D., de Groot, A. F. H., Wolf, K., Shi, L., Wei, T. Y., Durr, B. R., Chen, N. J., Vornbaumen, T., Wichmann, N. O., Mahamdeh, M. S., Pooladanda, V., Matoba, Y., Kumar, S., Kim, E., Bouberhan, S., Oliva, E., Rueda, B. R., Soberman, R. J., Bardeesy, N., Liau, B. B., Lawrence, M., Stokes, M. P., Beausoleil, S. A. & Bar-Peled, L. Systematic identification of anticancer drug targets reveals a nucleus-to-mitochondria pathway. Cell 186, 2361–2379 (2023).

36 Snyder, R. M., Mirabelli, C. K. & Crooke, S. T. Cellular association, intracellular distribution, and efflux of auranofin via sequential ligand exchange reactions. Biochem. Pharmacol. 35, 923–932 (1986).

37 Coffer, M. T., Shaw, C. F., 3rd, Hormann, A. L., Mirabelli, C. K. & Crooke, S. T. Thiol competition for Et3PAuS-albumin: a nonenzymatic mechanism for Et3PO formation. J. Inorg. Biochem. 30, 177–187 (1987).

38 Roberts, J. R., Xiao, J., Schliesman, B., Parsons, D. J. & Shaw, C. F., 3rd. Kinetics and Mechanism of the Reaction between Serum Albumin and Auranofin (and Its Isopropyl Analogue) in Vitro. Inorg. Chem. 35, 424–433 (1996).

39 Simon, T. M., Kunishima, D. H., Vibert, G. J. & Lorber, A. Screening trial with the coordinated gold compound auranofin using mouse lymphocyte leukemia P388. Cancer Res. 41, 94–97 (1981).

40 Mirabelli, C. K., Johnson, R. K., Sung, C. M., Faucette, L., Muirhead, K. & Crooke, S. T. Evaluation of the in vivo antitumor activity and in vitro cytotoxic properties of auranofin, a coordinated gold compound, in murine tumor models. Cancer Res. 45, 32–39 (1985).

41 Saba, N. S., Ghias, M., Manepalli, R., Schorno, K., Weir, S., Austin, C., Maddocks, K., Byrd, J. C., Kambhampati, S., Bhalla, K. & Wiestner, A. Auranofin Induces a Reversible In-Vivo Stress Response That Correlates With a Transient Clinical Effect In Patients With Chronic Lymphocytic Leukemia. Blood 122, 3819–3819 (2013).

42 Halatsch, M. E., Kast, R. E., Dwucet, A., Hlavac, M., Heiland, T., Westhoff, I. M. A., Debatin, K. M., Wirtz, C. R., Siegelin, M. D. & Karpel-Massler, G. Bcl-2/Bcl-xL inhibition predominantly synergistically enhances the anti-neoplastic activity of a low-dose CUSP9 repurposed drug regime against glioblastoma. Br. J. Pharmacol. 176, 3681–3694 (2019).

43 Halatsch, M. E., Kast, R. E., Karpel-Massler, G., Mayer, B., Zolk, O., Schmitz, B., Scheuerle, A., Maier, L., Bullinger, L., Mayer-Steinacker, R., Schmidt, C., Zeiler, K., Elshaer, Z., Panther, P., Schmelzle, B., Hallmen, A., Dwucet, A., Siegelin, M. D., Westhoff, M. A., Beckers, K., Bouche, G. & Heiland, T. A phase Ib/IIa trial of 9 repurposed drugs combined with temozolomide for the treatment of recurrent glioblastoma: CUSP9v3. Neurooncol. Adv. 3, vdab075 (2021).

44 Skrott, Z., Mistrik, M., Andersen, K. K., Friis, S., Majera, D., Gursky, J., Ozdian, T., Bartkova, J., Turi, Z., Moudry, P., Kraus, M., Michalova, M., Vaclavkova, J., Dzubak, P., Vrobel, I., Pouckova, P., Sedlacek, J., Miklovicova, A., Kutt, A., Li, J., Mattova, J., Driessen, C., Dou, Q. P., Olsen, J., Hajduch, M., Cvek, B., Deshaies, R. J. & Bartek, J. Alcohol-abuse drug disulfiram targets cancer via p97 segregase adaptor NPL4. Nature 552, 194–199 (2017).

45 Bindoli, A., Rigobello, M. P., Scutari, G., Gabbiani, C., Casini, A. & Messori, L. Thioredoxin reductase: A target for gold compounds acting as potential anticancer drugs. Coord. Chem. Rev. 253, 1692–1707 (2009).

46 Stafford, W. C., Peng, X. X., Olofsson, M. H., Zhang, X. N., Luci, D. K., Lu, L., Cheng, Q., Tresaugues, L., Dexheimer, T. S., Coussens, N. P., Augsten, M., Ahlzen, H. S. M., Orwar, O., Ostman, A., Stone-Elander, S., Maloney, D. J., Jadhav, A., Simeonov, A., Linder, S. & Arner, E. S. J. Irreversible inhibition of cytosolic thioredoxin reductase 1 as a mechanistic basis for anticancer therapy. Sci. Transl. Med. 10, eaaf7444 (2018).

47 Casini, A., Wai-Yin Sun, R. & Ott, I. MEDICINAL CHEMISTRY OF GOLD ANTICANCER METALLODRUGS in Metallo-Drugs: Development and Action of Anticancer Agents Ch. 7 (De Gruyter, 2018).

48 Yan, X., Zhang, X. S., Wang, L., Zhang, R., Pu, X. X., Wu, S. H., Li, L., Tong, P., Wang, J., Meng, Q. H., Jensen, V. B., Girard, L., Minna, J. D., Roth, J. A., Swisher, S. G., Heymach, J. V. & Fang, B. L. Inhibition of Thioredoxin/Thioredoxin Reductase Induces Synthetic Lethality in Lung Cancers with Compromised Glutathione Homeostasis. Cancer Res. 79, 125–132 (2019).

49 Raninga, P. V., Lee, A. C., Sinha, D., Shih, Y. Y., Mittal, D., Makhale, A., Bain, A. L., Nanayakarra, D., Tonissen, K. F., Kalimutho, M. & Khanna, K. K. Therapeutic cooperation between auranofin, a thioredoxin reductase inhibitor and anti-PD-L1 antibody for treatment of triple-negative breast cancer. Int. J. Cancer 146, 123–136 (2020).

50 Yin, N., Liu, Y., Weems, C., Shreeder, B., Lou, Y., Knutson, K. L., Murray, N. R. & Fields, A. P. Protein kinase Cι mediates immunosuppression in lung adenocarcinoma. Sci. Transl. Med. 14, eabq5931 (2022).

51 Chen, S. Y., Chiu, C. C., Hung, C. T., Pan, W. H., Chen, Y. C., Bow, Y. D., Li, W. J., Hsu, S. K., Lin, I. L., Wen, Z. H. & Wu, C. Y. Diphenyl disulfide potentiates the apoptosis of breast cancer cells through Bax proteolytic activation with accompanying autophagy. Environ. Toxicol. 38, 2022–2030 (2023).

52 Sibon, O. C. M. & Strauss, E. Coenzyme A: to make it or uptake it? Nat. Rev. Mol. Cell Biol. 17, 605–606 (2016).

53 McRae, M. P. Treatment of hyperlipoproteinemia with pantethine: A review and analysis of efficacy and tolerability. Nutr. Res. 25, 319–333 (2005).

54 Srinivasan, B., Baratashvili, M., van der Zwaag, M., Kanon, B., Colombelli, C., Lambrechts, R. A., Schaap, O., Nollen, E. A., Podgorsek, A., Kosec, G., Petkovic, H., Hayflick, S., Tiranti, V., Reijngoud, D. J., Grzeschik, N. A. & Sibon, O. C. Extracellular 4’-phosphopantetheine is a source for intracellular coenzyme A synthesis. Nat. Chem. Biol. 11, 784–792 (2015).

55 Gottlieb, N. L. Pharmacology of auranofin: overview and update. Scand. J. Rheumatol. Suppl. 63, 19–28 (1986).

56 Debnath, A., Parsonage, D., Andrade, R. M., He, C., Cobo, E. R., Hirata, K., Chen, S., Garcia-Rivera, G., Orozco, E., Martinez, M. B., Gunatilleke, S. S., Barrios, A. M., Arkin, M. R., Poole, L. B., McKerrow, J. H. & Reed, S. L. A high-throughput drug screen for Entamoeba histolytica identifies a new lead and target. Nat. Med. 18, 956–960 (2012).

57 Blocka, K. Auranofin versus injectable gold: Comparison of pharmacokinetic properties. Am. J. Med. 75, 114–122 (1983).

58 Walz, D. T., DiMartino, M. J., Griswold, D. E., Intoccia, A. P. & Flanagan, T. L. Biologic actions and pharmacokinetic studies of auranofin. Am. J. Med. 75, 90–108 (1983).

59 Shaw, C. F. Gold-based therapeutic agents. Chem. Rev. 99, 2589–2600 (1999).

60 Solmonson, A. & DeBerardinis, R. J. Lipoic acid metabolism and mitochondrial redox regulation. J. Biol. Chem. 293, 7522–7530 (2018).

61 Tsvetkov, P., Coy, S., Petrova, B., Dreishpoon, M., Verma, A., Abdusamad, M., Rossen, J., Joesch-Cohen, L., Humeidi, R., Spangler, R. D., Eaton, J. K., Frenkel, E., Kocak, M., Corsello, S. M., Lutsenko, S., Kanarek, N., Santagata, S. & Golub, T. R. Copper induces cell death by targeting lipoylated TCA cycle proteins. Science 375, 1254–1261 (2022).

62 Smith, A. R., Shenvi, S. V., Widlansky, M., Suh, J. H. & Hagen, T. M. Lipoic acid as a potential therapy for chronic diseases associated with oxidative stress. Curr. Med. Chem. 11, 1135–1146 (2004).

63 Steinbrueck, A., Sedgwick, A. C., Brewster, J. T., Yan, K. C., Shang, Y., Knoll, D. M., Vargas-Zuniga, G. I., He, X. P., Tian, H. & Sessler, J. L. Transition metal chelators, pro-chelators, and ionophores as small molecule cancer chemotherapeutic agents. Chem. Soc. Rev. 49, 3726–3747 (2020).

64 Holohan, C., Van Schaeybroeck, S., Longley, D. B. & Johnston, P. G. Cancer drug resistance: an evolving paradigm. Nat. Rev. Cancer 13, 714–726 (2013).

65 Ertl, P., Altmann, E. & McKenna, J. M. The Most Common Functional Groups in Bioactive Molecules and How Their Popularity Has Evolved over Time. J. Med. Chem. 63, 8408–8418 (2020).

66 Meyers, R. M., Bryan, J. G., McFarland, J. M., Weir, B. A., Sizemore, A. E., Xu, H., Dharia, N. V., Montgomery, P. G., Cowley, G. S., Pantel, S., Goodale, A., Lee, Y., Ali, L. D., Jiang, G., Lubonja, R., Harrington, W. F., Strickland, M., Wu, T., Hawes, D. C., Zhivich, V. A., Wyatt, M. R., Kalani, Z., Chang, J. J., Okamoto, M., Stegmaier, K., Golub, T. R., Boehm, J. S., Vazquez, F., Root, D. E., Hahn, W. C. & Tsherniak, A. Computational correction of copy number effect improves specificity of CRISPR-Cas9 essentiality screens in cancer cells. Nat. Genet. 49, 1779–1784 (2017).

67 Cai, D. M., Wang, J. J., Gao, B., Li, J., Wu, F., Zou, J. X., Xu, J. Z., Jiang, Y. L., Zou, H. Y., Huang, Z. H., Borowsky, A. D., Bold, R. J., Lara, P. N., Li, J. J., Chen, X. B., Lam, K. S., To, K. F., Kung, H. J., Fiehn, O., Zhao, R. Q., Evans, R. M. & Chen, H. W. ROR gamma is a targetable master regulator of cholesterol biosynthesis in a cancer subtype. Nat. Commun. 10, 4621 (2019).

68 Tepperman, K., Finer, R., Donovan, S., Elder, R. C., Doi, J., Ratliff, D. & Ng, K. Intestinal uptake and metabolism of auranofin, a new oral gold-based antiarthritis drug. Science 225, 430–432 (1984).

69. Luo, Y. L., Cao, B., Zhong, M. J., Liu, M. Y., Xiong, X. L. & Zou, T. T. Organogold(III) Complexes Display Conditional Photoactivities: Evolving From Photodynamic into Photoactivated Chemotherapy in Response to O-2 Consumption for Robust Cancer Therapy. Angew. Chem. Int. Ed. 61, e202212689 (2022).

